# Low Magnetic Fields Stimulate Cardiac Mitochondrial Bioenergetics with a Bell-Shaped Response: Possibly Via a Radical Pair Mechanism

**DOI:** 10.1101/2025.01.06.631559

**Authors:** Gisela Beutner, Huoy-Jen Yuh, Ilan Goldenberg, Douglas C. Wallace, George A. Porter, Arthur J. Moss, Shey-Shing Sheu

**Affiliations:** Department of Pediatrics, School of Medicine and Dentistry, University of Rochester, 601 Elmwood Avenue, Rochester, NY 14642 USA; Center for Translational Medicine, Department of Medicine, Sidney Kimmel Medical College, Thomas Jefferson University, 1020 Locust Street, Philadelphia, PA 19107 USA; Department of Medicine, School of Medicine and Dentistry, University of Rochester, 601 Elmwood Avenue, Rochester, NY 14642 USA; Center for Mitochondrial and Epigenomic Medicine, Department of Pediatrics, Division of Human Genetics, Children’s Hospital of Philadelphia and the Perelman School of Medicine, University of Pennsylvania, 3401 Civic Center Blvd. Philadelphia, PA 19104 USA

**Keywords:** Low magnetic field, heart mitochondria bioenergetics, mitochondrial respiration, electron transport chain activity, radical pair formation, electron spin transition

## Abstract

Studies have shown that low magnetic fields (LMFs) of less than 1 × 10^−3^ Tesla (T) affect numerous biological events, including bacteria and plant growth, bird migration, and human brain activity. On a cellular level, LMFs affect ion channel activities, intracellular Ca^2+^ concentrations, and mitochondrial reactive oxygen species (mROS) generation. However, the mechanisms that could account for these effects are controversial. Here, we show that applying a static LMF, ranging from ∼2.7 × 10^−4^ to ∼1.9 × 10^−3^ T, to mitochondria isolated from adult rat hearts produced a bell-shaped increase in maximal respiration (Vmax) up to 40%. A similar LMF-induced increase in Vmax was also observed in mitochondria isolated from rat hearts subjected to ischemia-reperfusion (I-R) injury. We then obtained data showing that LMF- mediated bell-shaped response was also observed in the activity of several enzymes involved in oxidative phosphorylation (OXPHOS), including Complexes II, III, and V, and citrate synthase. By contrast, similar LMF caused little change in the enzymatic activity of Complex I. Interestingly, mROS generation responded to LMF with an inverted bell-shaped decrease. We propose a radical pair mechanism of magnetoreception in cytochromes, catalytical reactions, and iron-sulfur clusters within the OXPHOS enzymes to explain how an LMF can increase the likelihood of electron spin transitions from singlet to triplet state and reverse it as the magnetic field strength further increases, resulting in a bell-shaped response. Our results indicate that a narrow range of LMF can enhance mitochondrial bioenergetics and decrease mROS. This may provide a non-invasive approach to treating diseases, such as I-R injury, when energy generation is compromised and oxidative stress is magnified.

## Introduction

Investigations on the effect of magnetic or electromagnetic fields (MFs or EMFs) applied to whole animals, organs, or cells have been widely reported, as summarized by Zadeh-Haghighi, et. al. (Zadeh-Haghighi and Simon 2022). The results reported are generally controversial, mainly due to the differences in the applied fields’ intensities, frequencies, and durations.

Studies of very low MFs (LMFs) on chemical reactions and biological activities in the last few decades have uncovered some potent effects (Greenebaum 2018, M. Sarraf 2020, Saletnik, Saletnik et al. 2022). At an LMF of less than 1 × 10^−3^ Tesla (T), it can either enhance or diminish the rates of specific chemical reactions, then reverse its effect when the LMF is increased further. This bell-shaped effect has been classified as a Low Field Effect (LFE) (Ulrich 1989, Barnes and Greenebaum 2015, Lewis, Fay et al. 2018). Three well-known examples of LFE in biological systems are migratory birds, plant growth, and bacteria quorum sensing, which orient and migrate by sensing the earth’s LMF of approximately 5 × 10^−5^ T (Xu, Jarocha et al. 2021, Xie 2022, Zhou, Tong et al. 2023). Quantum radical pair formation and electron spin transition have been hypothesized to explain LFEs (Y. Sakaguchi 1980, Ulrich 1989, Rodgers and Hore 2009).

Several reports have shown that LFE can affect mitochondrial functions. Human SH-SY5Y neuronal-like cells exposed to a uniform 2.2 × 10^−3^ T LMF for 24 hours resulted in a decreased mitochondrial membrane potential of up to 30% in association with increased reactive oxygen species (ROS) levels (Calabro, Condello et al. 2013). A square waveform LMF of 5 × 10^−3^ T at 50 Hz can increase mitochondrial activity and sperm mobility (Iorio, Delle Monache et al. 2011). Planarian growth can be impaired between 1 and 4 × 10^−4^ T, but its growth increases again at 5 × 10^−4^ T, associated with ROS production (Van Huizen, Morton et al. 2019). Static LMFs can alter cell death and proliferation, mainly through modulation of Ca^2+^ influx in cells (Fanelli, Coppola et al. 1999, Tenuzzo, Vergallo et al. 2009).

Mitochondrial ATP production involves the activity of the tricarboxylic acid (TCA) cycle to generate the substrates NADH and FADH_2_, which are oxidized by the Complexes I and II in the electron transport chain (ETC). The redox reactions through ETC Complexes involve several protein subunits, including ubiquinones, semiquinones, cytochromes, and iron-sulfur clusters, which can form radical pairs similar to the migrating birds’ magneto-sensing system (Xu, Jarocha et al. 2021, Zhou, Tong et al. 2023).

Taken together, these studies prompted us to address the following two questions: 1) Are mitochondrial OXPHOS activity and associated mitochondrial ROS (mROS) production in cardiac muscle cells modulated by LMF? 2) If they are, what are the probable mechanisms for this modulation? To address these questions, we investigated the effects of a static LMF, in a range of zero to ∼2 × 10^−3^ T, on the respiration rate, ROS generation, and activities of Complexes I, II, III, V, and several TCA cycle enzymes of mitochondria isolated from rat hearts. Our results show that a static LMF caused a bell-shaped increase in the maximal respiration rate (Vmax), the activity of several enzymes of the ETC and TCA cycle, and an inverted bell-shaped decrease in ROS levels. We then applied quantum mechanics to explain the experimental results theoretically in terms of affecting radical pair formation, hyperfine interaction, and electron spin interconversion.

## Results

### Setup for applying LMF to the oxygraph chamber

LMF was generated by a pair of coils mounted in the water jacket of the oxygen measurement chamber of an oxygraph (Hansatech, PP Systems, Amesbury, MA) (Figure 1A). The coils were held in place by a pair of plastic spools positioned next to the two water jacket outlets, as shown in Figures 1A & 1B. Each coil consists of a 24 gauge copper wire that was wound in 134 loops around each of the two spools and extended out of the water jacket outlet to connect to a DC power supply for providing various intensities of currents (in milli Ampere, mA) to generate different magnitudes of LMF. The distance between the centers of the two coils was 2.5 cm. The diameter of the coils was 2.5 cm, and the coil width was 0.9 cm. Since the two coils were separated by a distance 2 times the coil radius so that the entire chamber could be exposed to the applied LMF, the LMF inside the chamber was not entirely uniform, unlike the standard Helmholtz coils, which have a coil distance equal to the radius of the coils. Instead, the LMF distribution showed two identical peaks at the center of each coil and a valley between the two coils. The LMF at a location x from the center of the two coils is predicted from the theoretical calculation with the formula: B_0_= (μ₀NIR²/2) [(1/ (R² + (d - x)²))^3/2^ + (R^2^ / (1 + (d + x)²))^3/2^]; where B_0_ is the total MF, μ₀ is the permeability constant of free space, N is the number of turns per coil, I is the current in each coil, R is the radius of each coil, and d is the distance between the centers of the coils (blue dotted curve in Figure 1B). To validate the theoretical calculation, we experimentally measured the intensity of LMF with a Gauss meter along the center of the chamber as the function of distance from the bottom to the top of the chamber, with applied currents (I) ranging between 0 and 400 mA. These measured LMFs include the earth’s tiny LMF of 0.25-0.65×10^−4^ T. The results are shown in Figure 1C. The magnitudes of LMF with various intensities of the applied current all showed typical two peaked curves with the lowest valley values, approximately 60-70% of the peak values in 300 mA and 400 mA. This difference became smaller at the current ranges between 100-150 mA (green box in Figure 1C), which exerted the most significant effects on mitochondria (as described below in Figures 3-7).

**Figure 1:**
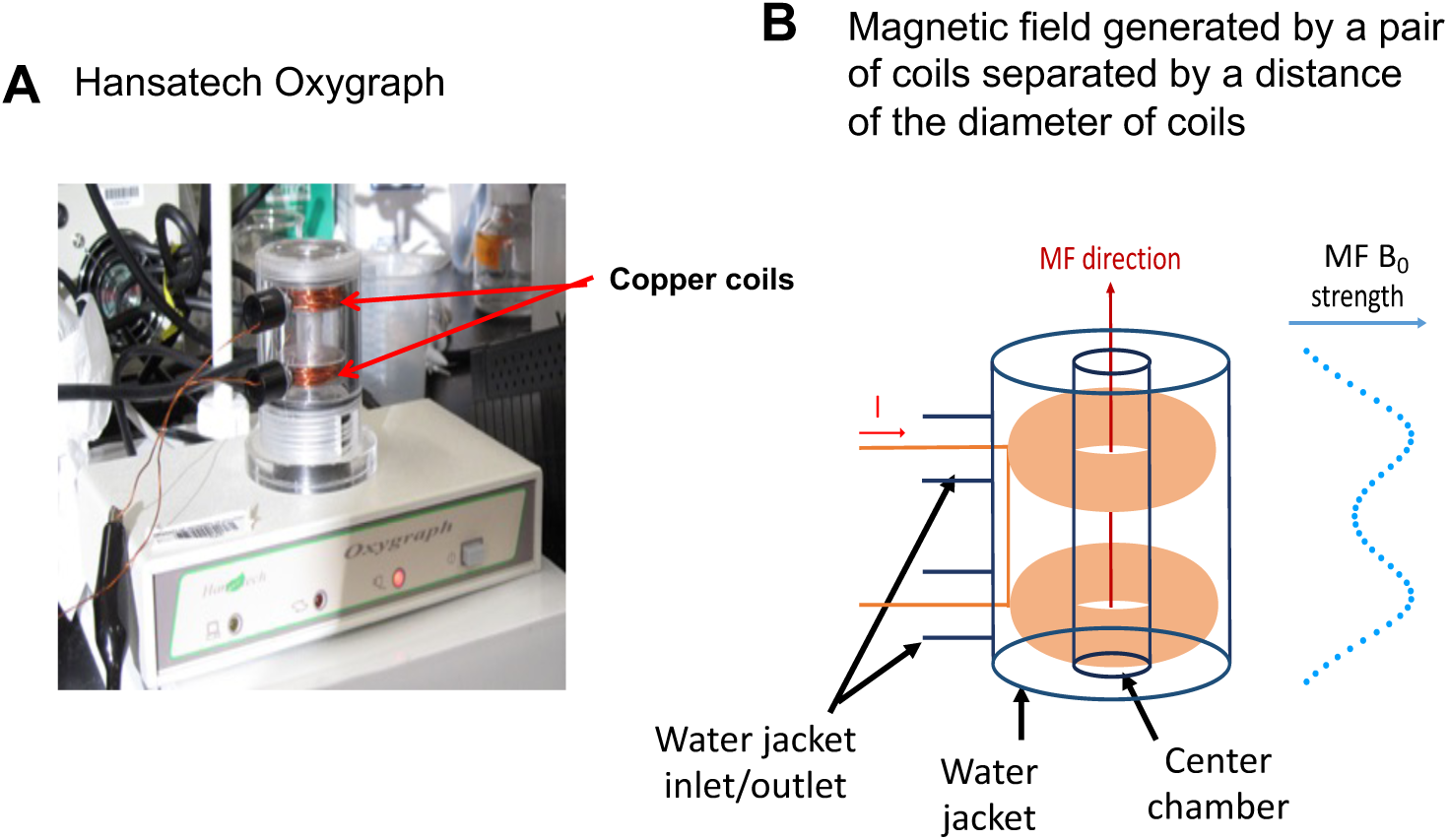
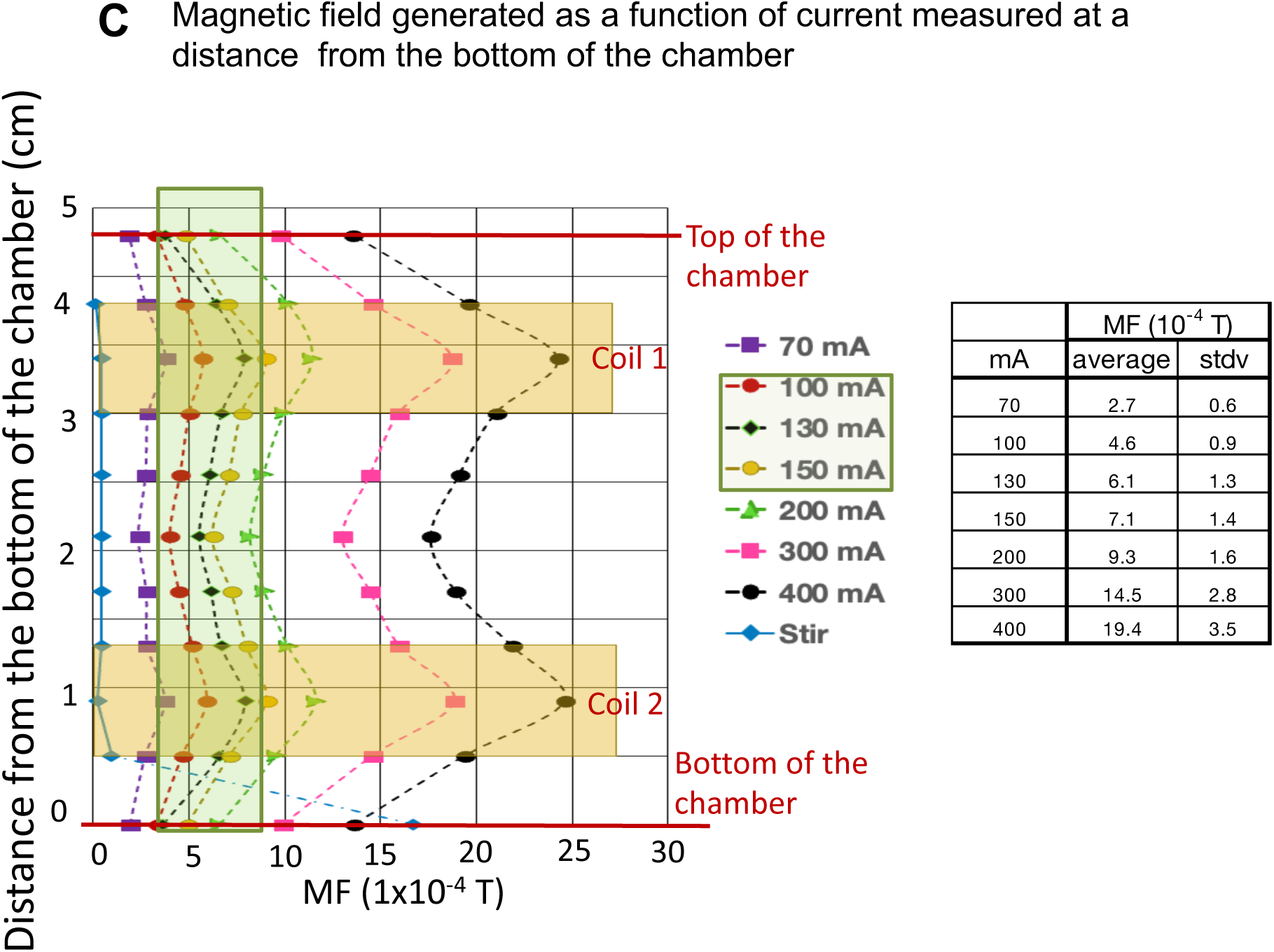
Experimental setup for measuring LMF effects on mitochondrial respiration. **A:** The water jacket of a Hansatech oxygraphy was retrofitted with two parallel copper coils mounted on plastic spools, which were connected through the inlet and outlet of the water jacket to a power supply. **B:** A schematic drawing of the coils (orange color). The average coil radius was 1.25 cm. The distance between the center of the two coils was 2.5 cm, which ensures that the bulk part of the chamber is exposed to the higher levels of MF generated. The ends of each coil’s copper wire are connected to a power supply to apply current (I) to the coils. The generated MF direction (red arrow) by applying current (I) and the theoretically calculated MF (B_0_) strength profile (blue dot curve) were also shown in the graph. **C:** MF intensities were measured with a Gauss meter every 0.5 cm from the bottom to the top inside the measuring chamber. MF generated by a magnetic stir bar was also measured (blue diamond). The green box shows the ranges of MF (100-150 mA) that produce the most significant effects on mitochondrial O_2_ consumption. The table on the right of Figure 1C shows the averaged MF magnitude of each applied I by averaging the measured MF from each location (data points in each connected curve).

To evenly disperse the isolated mitochondria in the oxygraph chamber with buffered solution, a small magnetic stir bar sitting at the bottom of the chamber was spun constantly by an oscillating magnet in the control unit of the oxygraph system directly below the measuring chamber. Figure 1C also shows the measurements of the intensity of the MF generated by this magnetic stirring bar spinning at various speeds, 50-75 % of the 900 rpm (blue diamond trace). LMF was ∼16×10^−4^ T at the bottom of the chamber and dropped steeply to 0.2-0.4 × 10^−4^ T at 0.4 cm above the bottom. This is approximately 1.5 to 10% of the highest (400 mA) and lowest (70 mA) fields applied for the experiments. Due to the contributions of the earth’s and stir bar’s MF, albeit small, during our experimental recordings, we present the LMF-mediated effects as a function of the applied currents (in mA). However, as a reference, the Table in Figure 1C shows the averaged LMF magnitude inside the chamber of those currents applied in our experiments.

Other control experiments validated that LMF did not affect oxygen concentrations of H_2_O, buffer solution, or mitochondria in a buffer solution without any substrates for ETC activation. Figures 2A-C show that the oxygen consumption rate remained constant upon alternating application of LMF (0, 100, 150, or 200 mA). Furthermore, using a mitochondria-free assay, we measured whether LMF alters the activity of the glucose oxidase, which oxidizes glucose into gluconic acid while reducing oxygen to hydrogen peroxide. The addition of 5 units of glucose oxidase to oxidize 10 mM glucose was not affected by the LMF generated by 70-300 mA (n=5; Figure 2D). Even though LMF generated by more than 200 mA tended to decrease glucose oxidase activity, this difference was insignificant (p≥0.05, n=4, ANOVA).

**Figure 2.**
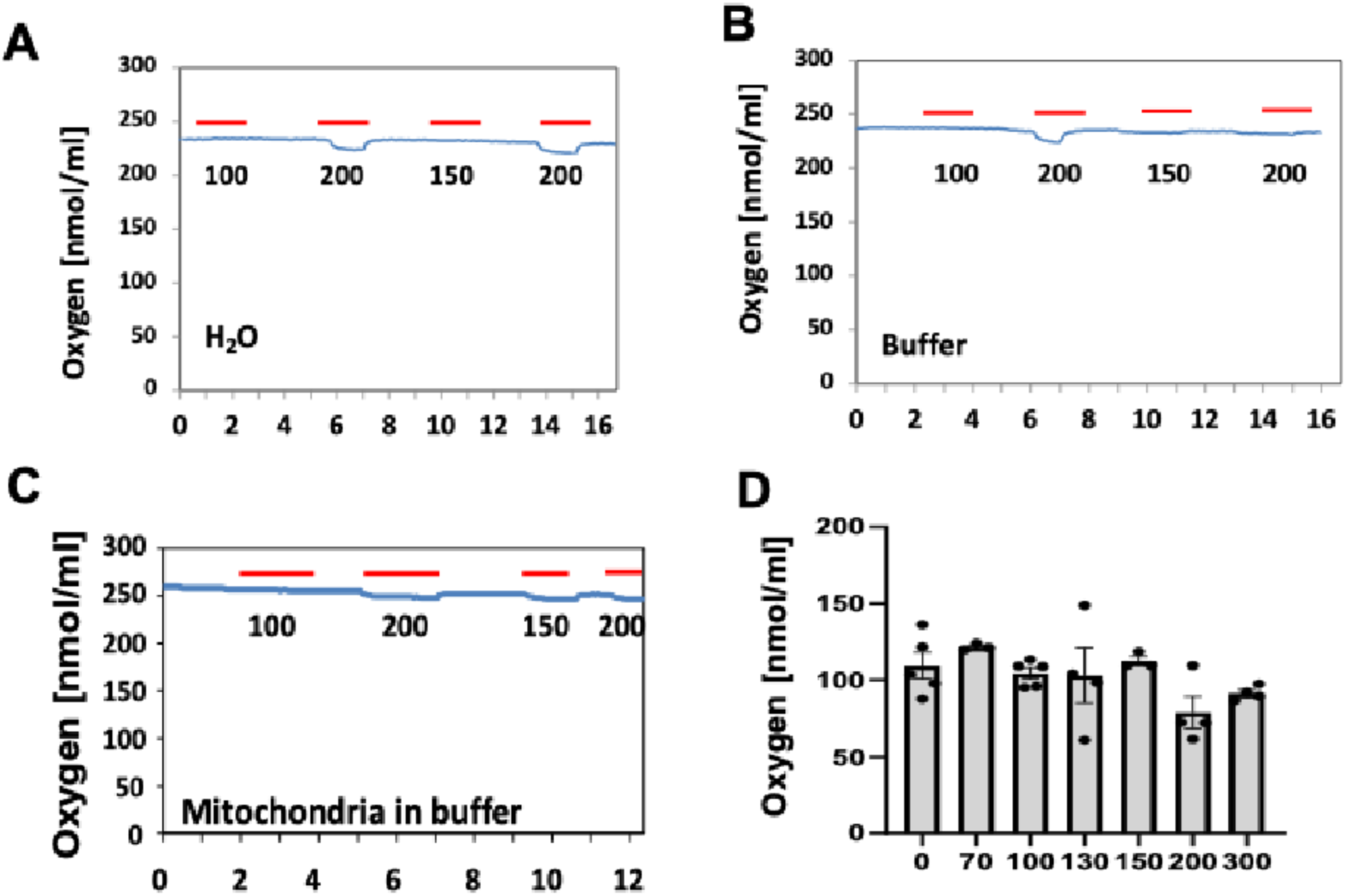
Lack of effects of LMF on oxygen concentrations of various control conditions. Experiments show LMF did not affect the oxygen concentrations of H_2_O (**A**), respiration buffer (**B**), mitochondria in respiration buffer without substrates (**C**), or glucose oxidase (**D**). In **A-C**, LMF generated by 100-200 mA (red bars) was alternated with no field. In **D:** LMF from 70-300 mA did not affect oxygen consumption by a glucose oxidase reaction.

### LMF increases Vmax and mitochondrial coupling

OXPHOS is the most important mechanism for generating cellular energy in the form of ATP and can be assessed by measuring mitochondrial oxygen consumption. Therefore, the first set of experiments aimed to determine whether LMF affects mitochondrial oxygen consumption. The blue line graph at the bottom panel of Figure 3A shows the time course of experimental manipulations for assessing the effects of LMF on oxygen consumption of isolated rat heart mitochondria (see figure legend for detailed description). The top panel of Figure 3A shows a representative oxygraph recording of V_0_ (indicated by the addition of malate and glutamate, MG) and Vmax (indicated by the addition of ADP) in the absence (black trace) and the presence of a 100 mA generated LMF (red trace). The LMF did not alter the slope of oxygen consumption of V_0_ but increased the slope of oxygen consumption from 26.74 O_2_/min to 40.36 O_2_/min (51% increase) at Vmax. The effects of a range of LMFs (0-200 mA) on V_0_ and Vmax are shown in Figures 3B and 3C, respectively. The data show that LMFs did not affect V_0_ but caused a bell-shaped biphasic increase in Vmax with a peak response at 100 mA (∼4.6 × 10^−4^T). These data points were then used to calculate the respiratory control ratio (RCR, Vmax/V_0_), a measure of mitochondrial coupling (Figure 3D). In the presence of 100 mA, RCR increased significantly from 3.38 ± 0.185 (control) to 4.83 ± 0.287 (43% increase, n=7, p ≤ 0.05, ANOVA, Figure 3D, 100 mA data point). The effects of LMFs (0-200 mA) on RCR showed a non-linear correlation (bell-shaped) to the applied LMF and peaked at around 100 mA (∼4.6 × 10^−4^T) (Figures 3C & D, respectively). Non-linear polynomial regression analysis shows a second-order best fit for Vmax and RCR response (Figure 3E).

**Figure 3.**
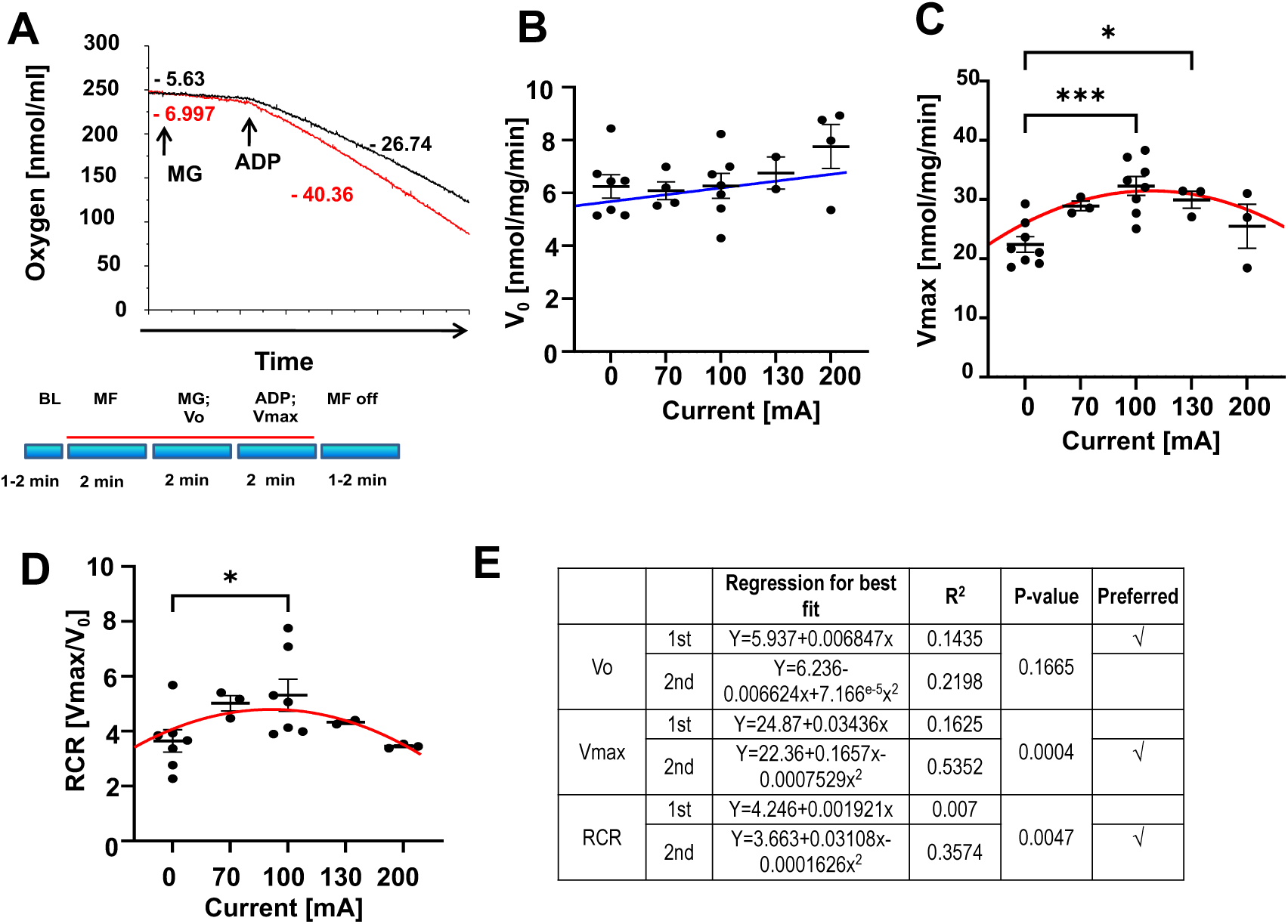
Mitochondrial respiration as a function of applied LMF. **A:** The blue bars at the bottom show the time course of experimental manipulations: Baseline (BL), the magnetic field applied (MF) throughout the time labeled by a red line, the addition of substrate (MG; 3 mM malate and 5 mM glutamate for initiating V_0_, and 1 mM ADP for initiating Vmax, and then switched off MF. The top graphic panel shows a representative recording of V_0_ and Vmax with and without LMF applied. Black trace and values: no MF; red trace and values: LMF generated by 100 mA. LMF significantly increased (51%) the rate (slope) of Vmax. **B**: V_0_ was not affected by LMF. **C**: Vmax increased by LMF with an asymmetric return bell-shaped curve. **D**: RCR (ratio of Vmax/V_0_) increased by LMF with an asymmetric return bell-shaped curve. **E**: Summary of non-linear regression of polynomial fittings with R^2^ and p-value. N=2-7; * p≤0.05; ***p≤0.005.

Ischemia-reperfusion (I-R) causes severe heart injury, assumed to be resulting from the burst of mROS generation upon O_2_ reperfusion (Chouchani, Pell et al. 2016). To investigate the effect of LMF on mitochondrial respiration following I-R, Langendorff perfused hearts were first exposed to the stop flow condition (“ischemia”) for 20 minutes and then reperfused with fully oxygenated solution (reperfusion) for 20 minutes. The hearts were then used for mitochondria isolation. For comparison, control hearts were perfused with oxygen-saturated buffer continuously for 40 minutes, and then the hearts were used for mitochondria isolation. LMF did not affect V_0_ in control and I-R-treated mitochondria (Figures 4A & 4E). Vmax in the absence of LMF was approximately 2-fold higher in control conditions than in I-R-treated mitochondria (Figures 4B & 4F, 0 mA data point, respectively), indicating some OXPHOS defects in I-R-treated mitochondria. However, LMFs induced similar bell-shaped % increases of Vmax, both in control and I-R-treated mitochondria, with a maximum increase approximately of 30 % compared to baseline values without LMF treatment (0 mA, set as 100%) (Figure 4D, n=3; p≤0.05). The relative increases in RCR by LMF were similar between control and I-R-treated mitochondria, as shown in Figures 4C, 4G, & 4H. The non-linear curve fitting of both experimental groups again shows an asymmetric 2-order bell-shaped effect by LMF on Vmax and RCR (Figure 4I Table).

**Figure 4.**
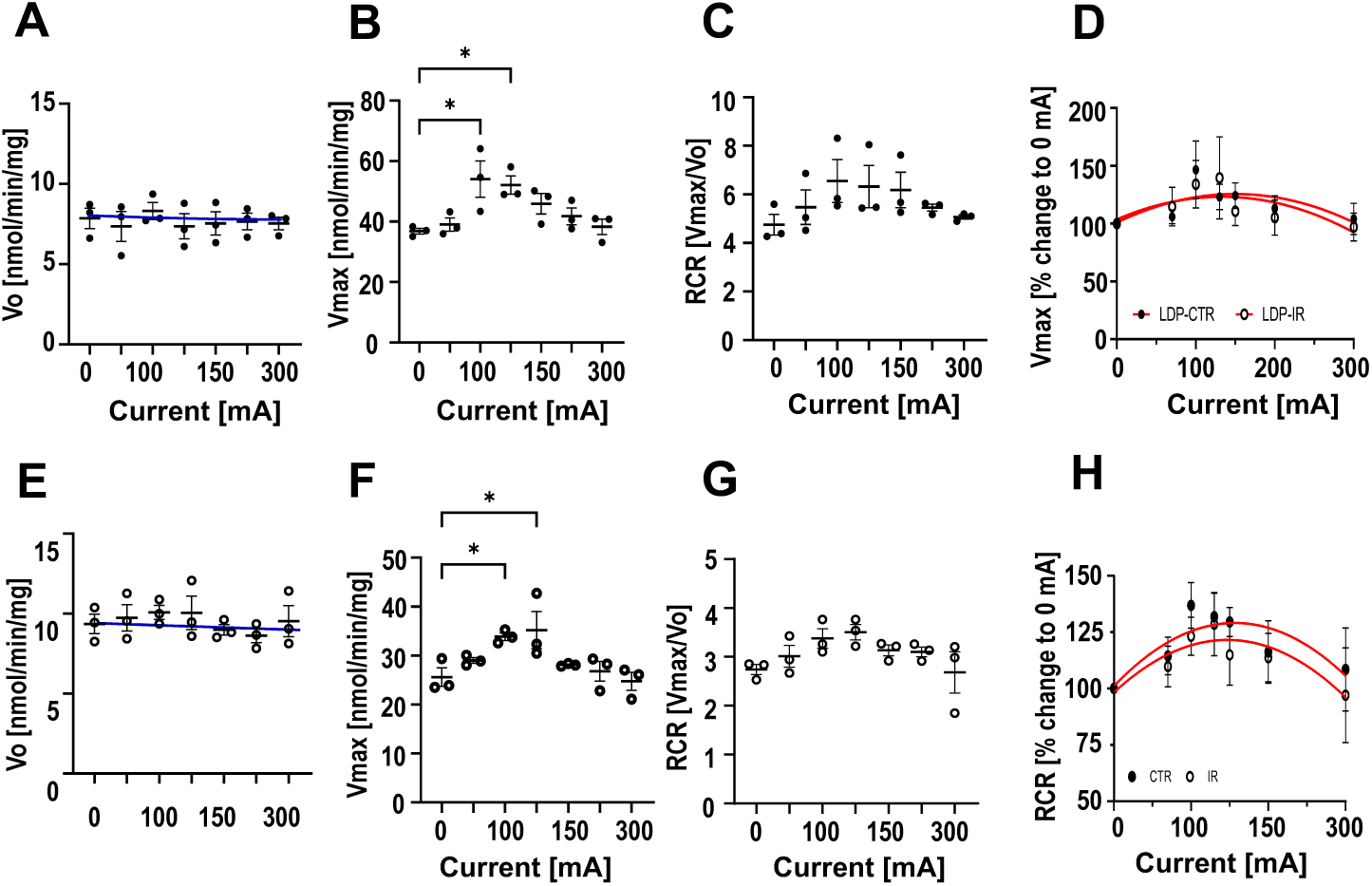
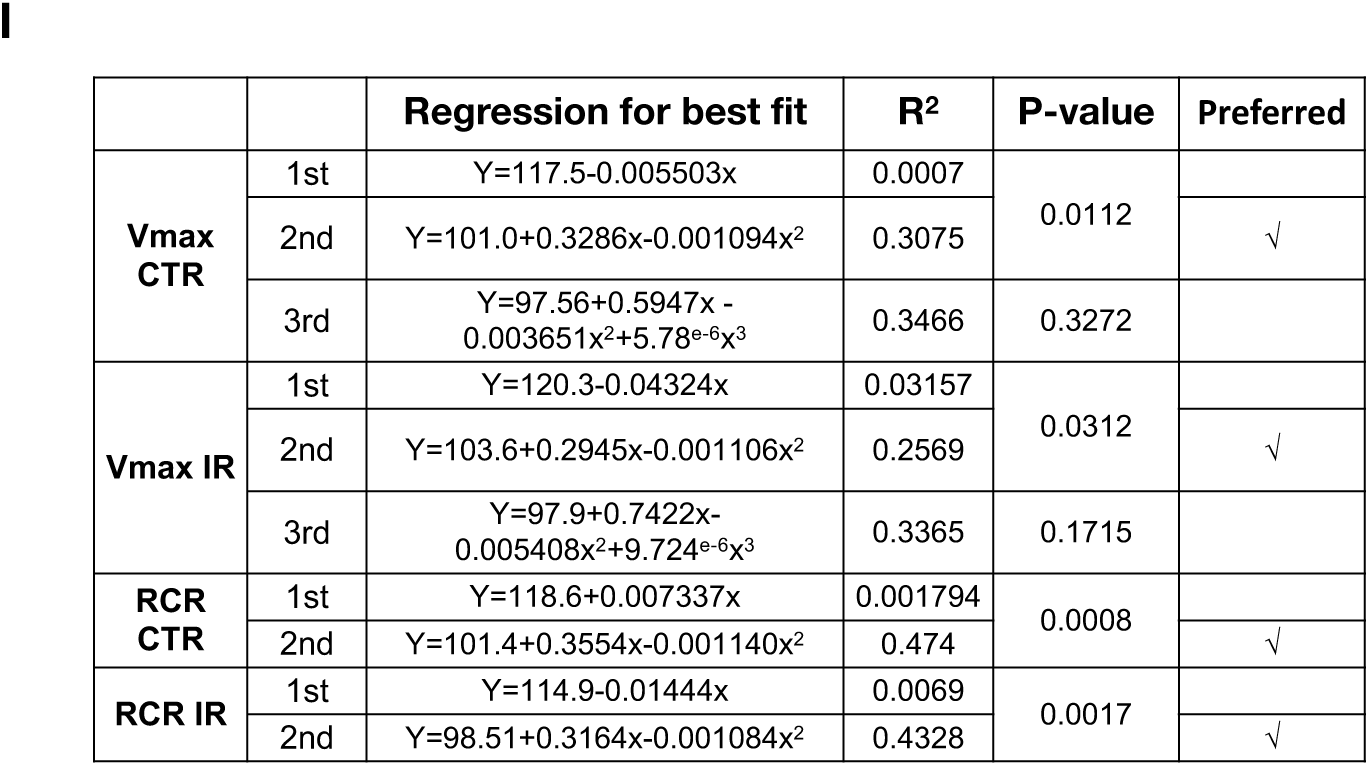
Oxygen consumption as a function of the applied LMF in mitochondria isolated from hearts after Langendorff perfusion (LDP), control (CTR, ●), and I-R (IR, ○). **A & E**: V_0_ in both preparations was not affected by LMF. Vmax was higher in control (**B**) than in I-R (**F**) in the absence of LMF. However, LMF induced a similar % increase in Vmax of both preparations (**B & F**) (**D**). RCR (Vmax/V_0_) was higher in the control (**C**) than in I-R (**G**), but the LMF produced a similar RCR increase in the control and I-R (**H**). Non-linear regression preferred a polynomial second-order fitting. N=3; * p≤0.05 (**I**).

### LMF increases the enzymatic activity of the Complexes II, III, and V of the ETC

Mitochondrial oxygen consumption is a readout for ETC activities. To determine which complexes of the mitochondrial ETC were responsible for increased oxygen consumption, we measured their activities after exposure to LMF. Mitochondria were first equilibrated in the respective assay mix (without inhibitors and starters) for 5 minutes at room temperature and then exposed to LMF for 2 minutes. Immediately after the exposure to LMF, inhibitors and starters were added, and the activity was recorded.

The Complex I activity was not affected by LMF (Figure 5A). A negative slope was shown in the linear regression for best fitting the data, suggesting a decreasing trend of the effect, but not statistically significant (p>0.05). The activities of Complexes II and III increased significantly (p≤0.01 and p≤0.05, respectively) by LMF, with an asymmetrical bell-shaped response that peaked around 100-150 mA, similar to that of oxygen consumption (Figures 5B, 5C, and 5E).

**Figure 5:**
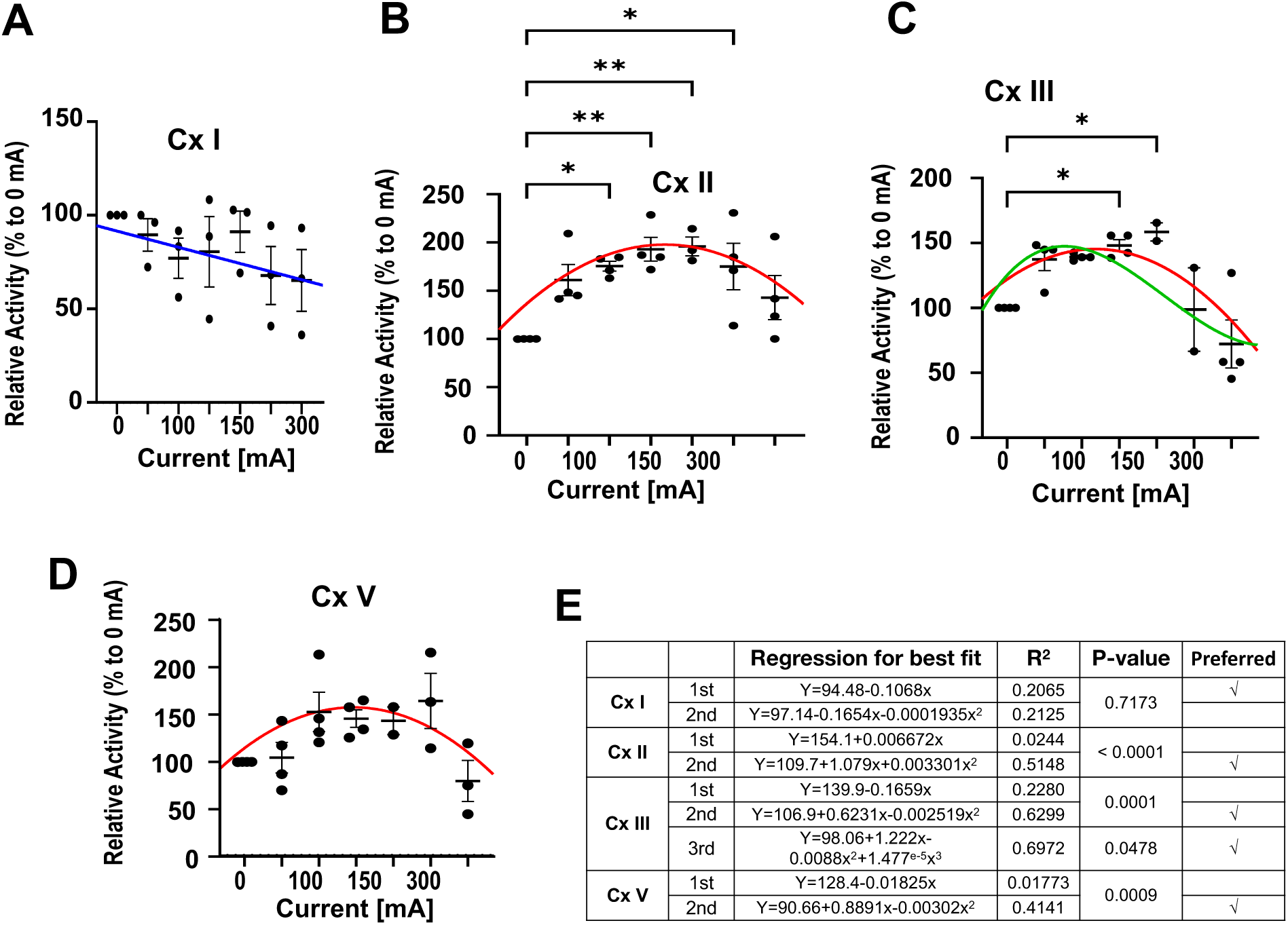
Relative enzymatic activity of the ETC Complexes I, II, III, and V in response to LMF. Activity was normalized to no LMF treatment (0 mA) and as 100 %. **A:** Complex I. **B:** Complex II. **C:** Complex III. **D:** Complex V. **E**: Table summarizing non-linear fitting: first (blue line in A), second (red line in B, C, D), and third (green line in C) order. N=2-4; * p≤0.05; **p≤0.01.

Complex V activity also showed an asymmetrical bell-shaped response after exposure to LMF, which peaked at around 130 mA (Figure 5D), but no significant differences statistically were apparent (p> 0.05).

Thus, LMF-mediated increases of the ETC Complexes II, III, and V could be the reason for an increase in Vmax and the RCR. This data set leads to whether the upstream of the ETC pathways also respond to LMF. Therefore, we measured the effect of LMF on several enzymes of the TCA cycle.

### LMF increases the activities of several TCA cycle enzymes

To determine if LMF also changed the activities of specific enzymes in the TCA cycle, we measured its effects on citrate synthase, succinate dehydrogenase (SDH, complex II), and malate dehydrogenase. Applying LMF showed a bell-shaped increase in citrate synthetase activity, peaking at ∼ 130 mA (Figure 6A). SDH is the only enzyme that participates in the citric acid cycle and the electron transport chain (Complex II). Applying LMF showed a bell-shaped increase of succinate dehydrogenase activity peaking at ∼130 mA (Figure 6B). LMF from 0-300 mA did not change malate dehydrogenase activity except for a single-point increase at 100 mA (Figure 6C). Regression analysis further confirmed a polynomial non-linear response for citrate synthase and succinate dehydrogenase but a linear response for malate dehydrogenase (Figure 6D).

**Figure 6:**
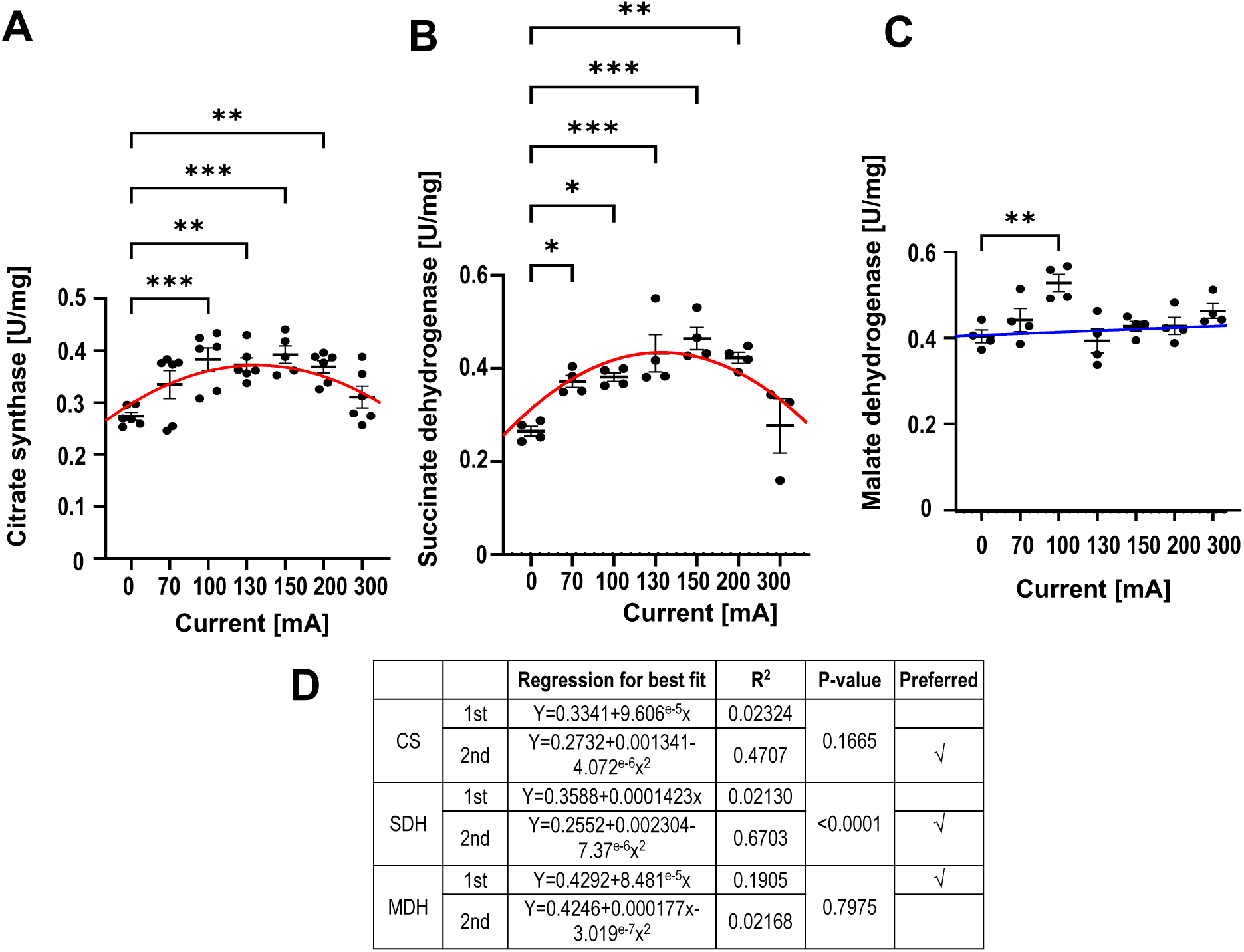
TCA cycle enzyme activity of citrate synthase, succinate, and malate dehydrogenases as a function of LMF and their non-linear fittings. **A:** Citrate synthase. **B:** Succinate dehydrogenase **C**: Malate dehydrogenase. **D**: The table summarizing the analysis of the fittings shows second-order best fit for citrate synthase and succinate dehydrogenase and first-order for malate dehydrogenase. N=3-6; * p≤0.05; **p≤0.01***p≤0.005.

### LMF decreases ROS levels during Vmax

To determine if LMF-mediated changes in mitochondrial respiration, the enzymatic activity of the ETC Complexes, and several TCA cycle enzymes were associated with an altered mROS production, we assayed the mitochondrial production of H_2_O_2_ in response to LMF. Mitochondrial ROS generation at conditions resembling V_0_ was unaffected by LMF (Figure 7A). However, at the conditions for assaying Vmax (M/G+ ADP), LMF caused an inverted bell-shaped response, dropping to a minimum at 100-130 mA (n=3-10; p ≤ 0.05; Figure 7B). Inhibition of Complex III with antimycin A (2 µg/ml) caused an increase in H_2_O_2_ independent of the applied LMF (Figure 7C). Regression analysis showed that a third-order polynomial best fitted the inverted response of H_2_O_2_ to LMF at Vmax (Figure 7D). This set of data indicates that an LMF not only increases Vmax but also limits the generation of ROS at the same time

**Figure 7:**
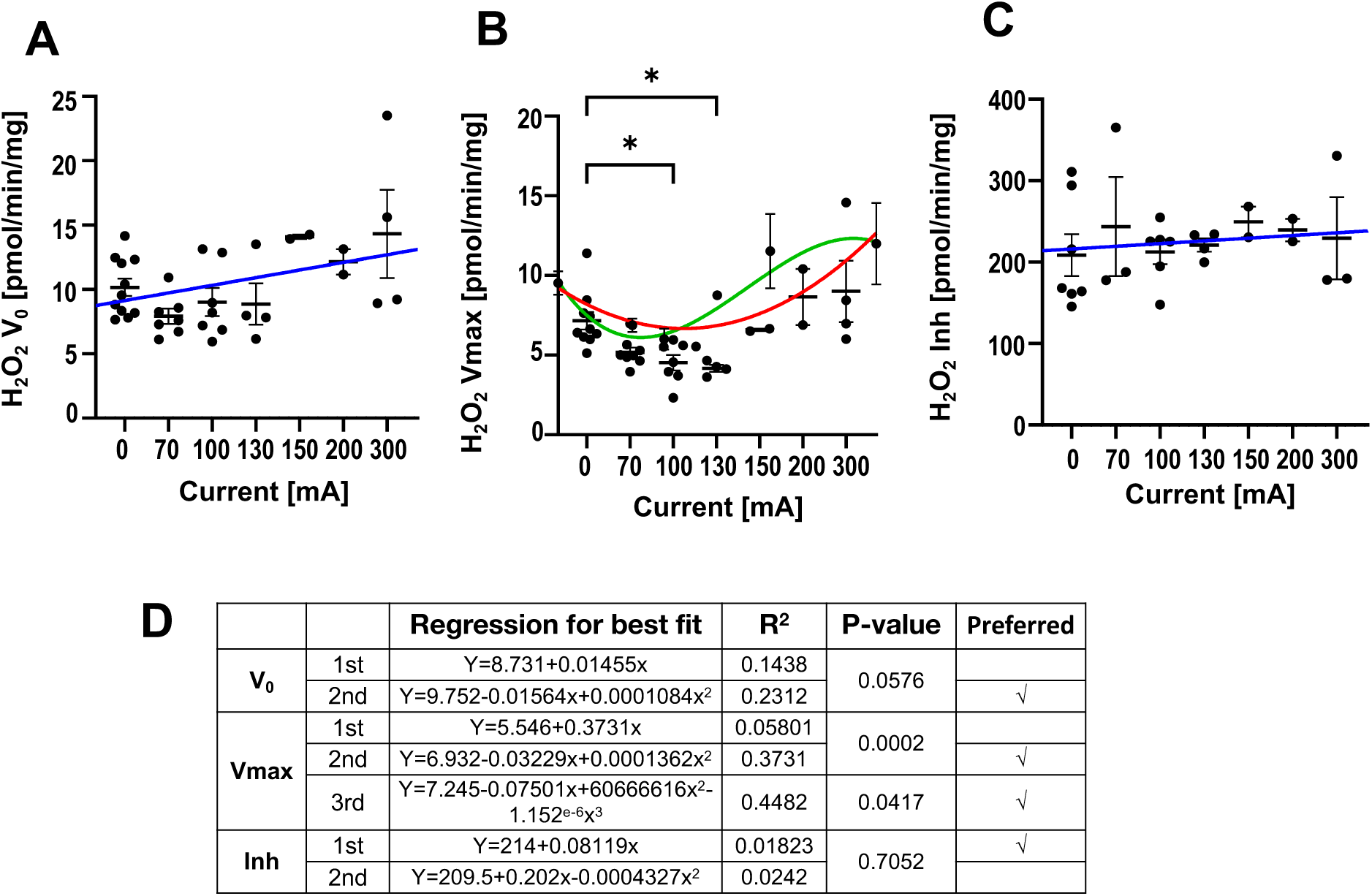
LMF modifies the generation of ROS. **A:** Mitochondrial ROS generation was unaffected in substrate-mediated respiration (V_0_). **B:** LMF decreases the generation of ROS in maximal respiring mitochondria (Vmax). **C:** The addition of Complex III inhibitor, antimycin A (inh, 2 µg/ml), increased basal ROS generation independently of applied LMF. **D:** Summary of non-linear fittings: blue first-order, red second-order, and green third-order fittings. N=2-11; * p≤0.05.

## Discussion

In this study, we observed the LFE in the cardiac mitochondria isolated from rats. Applying increasing strengths of an LMF from 0 to ∼14.5×10^−4^ T (0-300 mA) caused a striking bell-shaped increase of mitochondrial oxygen consumption, the activity of the Complexes II, III, and V, and the activity of the TCA cycle enzymes citrate synthase and succinate dehydrogenase.

The effects were detected between ∼2.7 and ∼9.3×10^−4^ T (70 and 200 mA, respectively). The maximum effects were observed between ∼4.6 to ∼6.1×10^−4^ T (100 to 130 mA). No effects were detected at ∼14.5 and ∼19.4×10^−4^ T (300 and 400 mA), which was the range that included the intensity of ∼16×10^−4^ T generated by the magnetic stir bar at the bottom of the chamber. These LFEs are in contrast to non-responsive or a modest linear decline (as suggested by curve fitting) in Complex I activity with increasing LMF and an inverted bell-shaped response curve of mitochondrial H_2_O_2_ production, which dropped to a minimum between ∼4.6 to ∼6.1×10^−4^ T. Others have previously reported increases in mitochondrial activity by LMF (Morabito, Rovetta et al. 2010, Dragicevic, Bradshaw et al. 2011, Iorio, Delle Monache et al. 2011, Zadeh-Haghighi and Simon 2022). Unique to the results in our studies is a non-linear bell-shaped response of several OXPHOS enzymes to a relatively narrow range of LMF. An intriguing question is: how can LMF generate strikingly similar responses among various OXPHOS proteins with very different chemical structures? In addition, what is the explanation for the inverted LMF responses between ROS and OXPHOS proteins? We attempt to address these two questions below.

### Redox reactions during OXPHOS

In the process of OXPHOS, as shown in Figure 8, Complex I oxidizes NADH to NAD^+^, and the electrons are transferred to a flavin mononucleotide (FMN) and then to a cascade of iron-sulfur (FeS) centers to exit Complex I by reducing coenzyme Q (CoQ) to CoQH_2_. Similarly, succinate is oxidized by succinate dehydrogenase (Complex II), with the electrons transferred to flavin adenine dinucleotide (FAD to FADH_2_) and via internal FeS center to CoQ. Electrons from CoQH_2_ are transferred to Complex III, harboring cytochromes b and c_1_, and Rieske FeS protein, and then shuttled through cytochrome c to Complex IV with cytochrome a, cytochrome a_3_, and two Cu centers. Complex IV transfers electrons from 4 cytochrome c molecules to molecular oxygen to produce two water molecules. As the electrons transverse Complexes I, III, and IV, protons are pumped from the mitochondrial matrix into the intermembrane space, creating a proton gradient that is used to drive the rotation of F_o_F_1_-ATP synthase, which allows the synthesis of ATP from ADP and Pi (inorganic phosphate).

**Figure 8:**
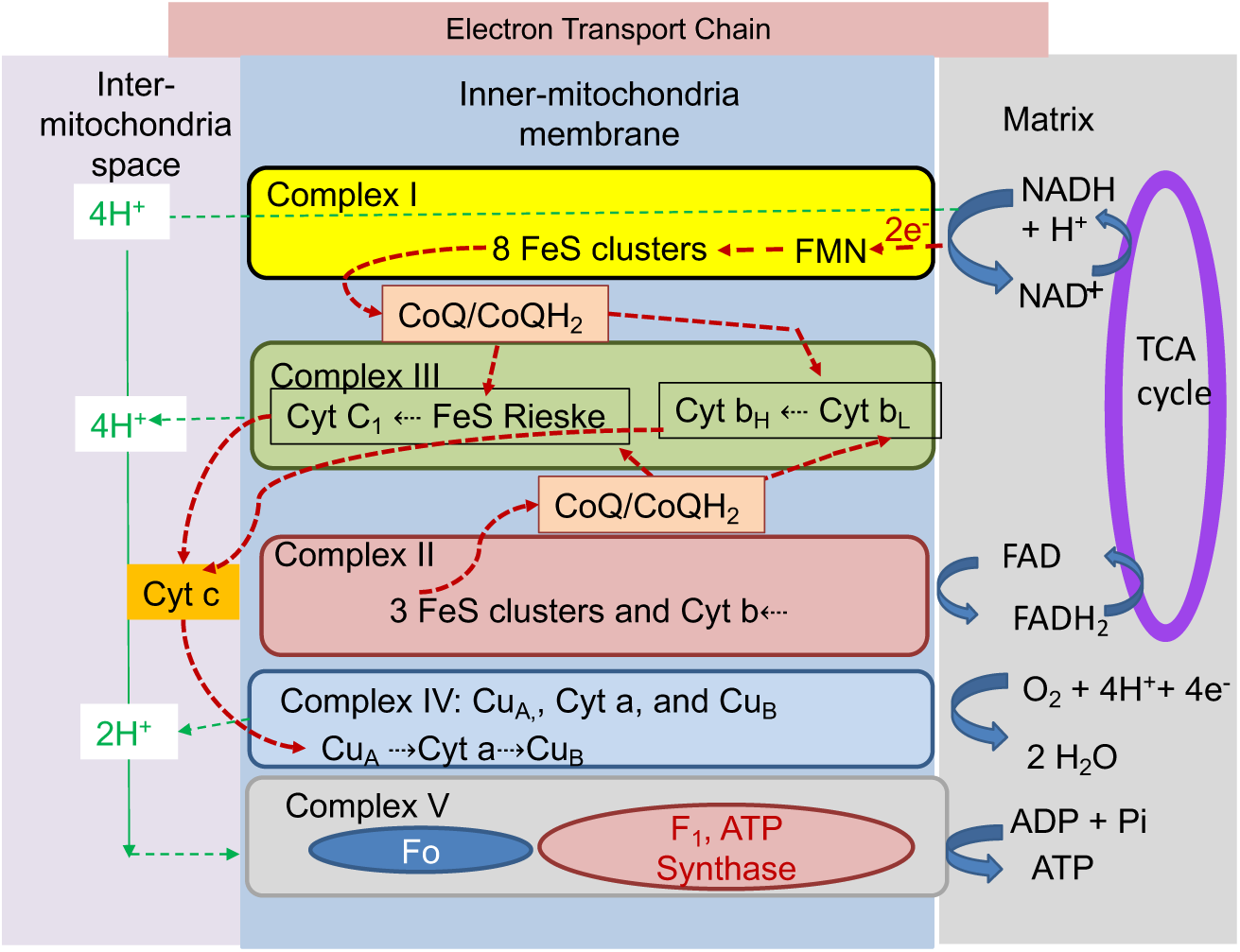
The schematic diagram of mitochondrial electron transport processes and ATP synthase.

mROS are thought to be generated primarily on Complexes I and III. In Complex I, they are generated mainly through the FMN via electron leak or electron backflow, with one electron transfer generating superoxide anion (O_2_^●-^) or a two-electron transfer to generate H_2_O_2_. Two O_2_^●-^ can also be dismutated by Manganese superoxide dismutase to generate H_2_O_2_.

Taken together, all these processes in OXPHOS involve electron transfer redox reactions. The present results, showing that an LMF can significantly affect the activity of OXPHOS-mediated electron transfer redox reactions, impose an intriguing question: what mechanism can account for this LFE in isolated cardiac mitochondria? To propose such a mechanism, it is essential first to describe the quantum mechanics of LMF influences on electron spin moments and radical pair reactions, as elaborated in the Supplementary Materials. Briefly, In quantum mechanics, electron spin refers to an electron’s intrinsic spin angular momentum. In a classic model, a single electron’s spin moment (m_s_) precesses in a cone shape along the MF with either up or down directions to the field (Supplementary Figure 1A). For those with two paired electrons that have opposite spin directions to each other, their spins cancel each other out, resulting in a net spin S being 0 (S_0_) with only one spin state of m_s_ = 0 (Supplementary Figure 1B). For those with two unpaired electrons, the net spin S=1 with three spin states of m_s_ of 0, +1, and −1 (Supplementary Figure 1B). Only electron spin moments of molecules or atoms can directly interact with an MF (Eason 2019). Non-zero electron spins can interact with an MF, known as the Zeeman effect, to alter their energy. In addition, the MF also alters electron spins’ Larmor procession speed. Together, these can affect the interconversion between different spin states (e.g., singlet and triplet states) and, as such, change the activities of electron transfer reactions.

Our results show that ∼5 to 6×10^−4^ T can increase Vmax up to 40%, which is surprising as the interaction energy with such an LMF is several orders of magnitude below the thermal energy of kT (Boltzmann constant x temperature), about 2-3×10^−3^ eV (electron volt) at room temperature, and the strength of any chemical bonds. Similar LMF-mediated effects have been detected in photochemical reactions when molecules absorb light energy to enter their excited state. An example of these LMF-induced photochemical reactions is the ability of migrating birds to use cryptochrome protein in the eye to sense Earth’s LMF for navigation (Qin, Yin et al. 2016, Xu, Jarocha et al. 2021, Zhou, Tong et al. 2023). To interpret these LFEs, a radical-pair mechanism involving the formation of radical pairs and electron spin interaction with the LMF has been proposed (see Supplementary Materials for a detailed description). The radical pair mechanism suggests that an external LMF enhances the transition of the singlet spin state (S_0_) to the triplet spin state (T_0_) of a radical pair intermediate A*B* in a chemical reaction before proceeding to the final product C (Supplementary Figures 2A & B). This increase in T_0_ is due to hyperfine suppression by the LMF. However, as the intensity of LMF increases, the Δg effect becomes dominant in decreasing T_0_. The summation of these two opposite effects results in a bell-shaped increase in the T_0_ population (Supplementary Figure 2C). Below, we attempt to apply this theory of LMF interaction with electron spin transition in the radical pair reaction to explain the potential mechanisms for our experimental results.

### Proposed mechanisms

Except for Complex I and malate dehydrogenase, the rest of this study’s ETC and TCA cycle proteins show a bell-shaped response to LMF, manifesting LFE attributed to the radical pair mechanism involving magnetically sensitive intermediate molecules (Ulrich 1989, Rodgers and Hore 2009, Mouritsen 2016, Lewis, Fay et al. 2018). Our results show a typical LFE starting with a radical pair in a singlet state. As discussed in Supplementary Materials, the spin-coupled radical pair can be two separate radicals loosely coupled or resulting from a charge separation to different locations within a molecule. The spatial separation of the two radicals is around 1 nm so that the energy levels of their singlet and triplet states are comparable to enable hyperfine disturbance by the external LMF. This poses an important question: why do these proteins involved in mitochondrial OXPHOS have similar capabilities to form radical pair intermediates under LMF exposure? To address this question, it is helpful to depict the steps of redox reactions of each ETC Complex, as described in Figure 8. Iron-sulfur clusters, which support the transport of electrons (Read, Bentley et al. 2021), are present in Complexes I, II, and III, while cytochromes are present in Complexes II, III, and IV. Complex IV contains 2 heme and 2 copper centers, and Complex V does not contain any of them but has F_o_F_1_ rotors driven by proton gradient to produce ATP. Can any of them form a “singlet” radical pair with characteristics described in Supplementary Materials?

Complex II and III contain cytochromes. All cytochromes (b, b_L_, b_H_, c_1_, and c) consist of iron centers, Fe^2+^/Fe^3+^, coordinated to the porphyrin derivatives, which have delocalized π orbitals (Figure 9) (Mowat 2013). Fe^2+^ has two paired orbital electrons at a singlet spin state. It transfers one electron to the following redox site by first separating one electron from itself, Fe^2+^(cyt) → Fe^3+^(cyt) + e^−^, where cyt is cytochrome coordinated porphyrin derivatives, and then transfers this electron to the following ETC site. The first electron-separating step is likely a rate-limiting step. The first separated electron could reside intermittently on the delocalized π orbitals before transferring to the following ETC site. This charge separation within a large molecule generates a spatially separated radical pair. Recent theoretical calculations have suggested that stabilized intermediate triplet states of Fe*^3+^-porphyrin*^1-^ with low-lying energy levels can potentially exist (Alavi 2018, Li Manni, Kats et al. 2019). The stabilization comes from the delocalization of electrons in the π orbitals of the coordinated porphyrin, as shown in Figure 9. It is reasonable to hypothesize that all cytochromes have a similar delocalization of low-lying triplet state intermediates, which can then go through a radical pair mechanism to generate LFE, as observed in Complexes II and III.

**Figure 9:**
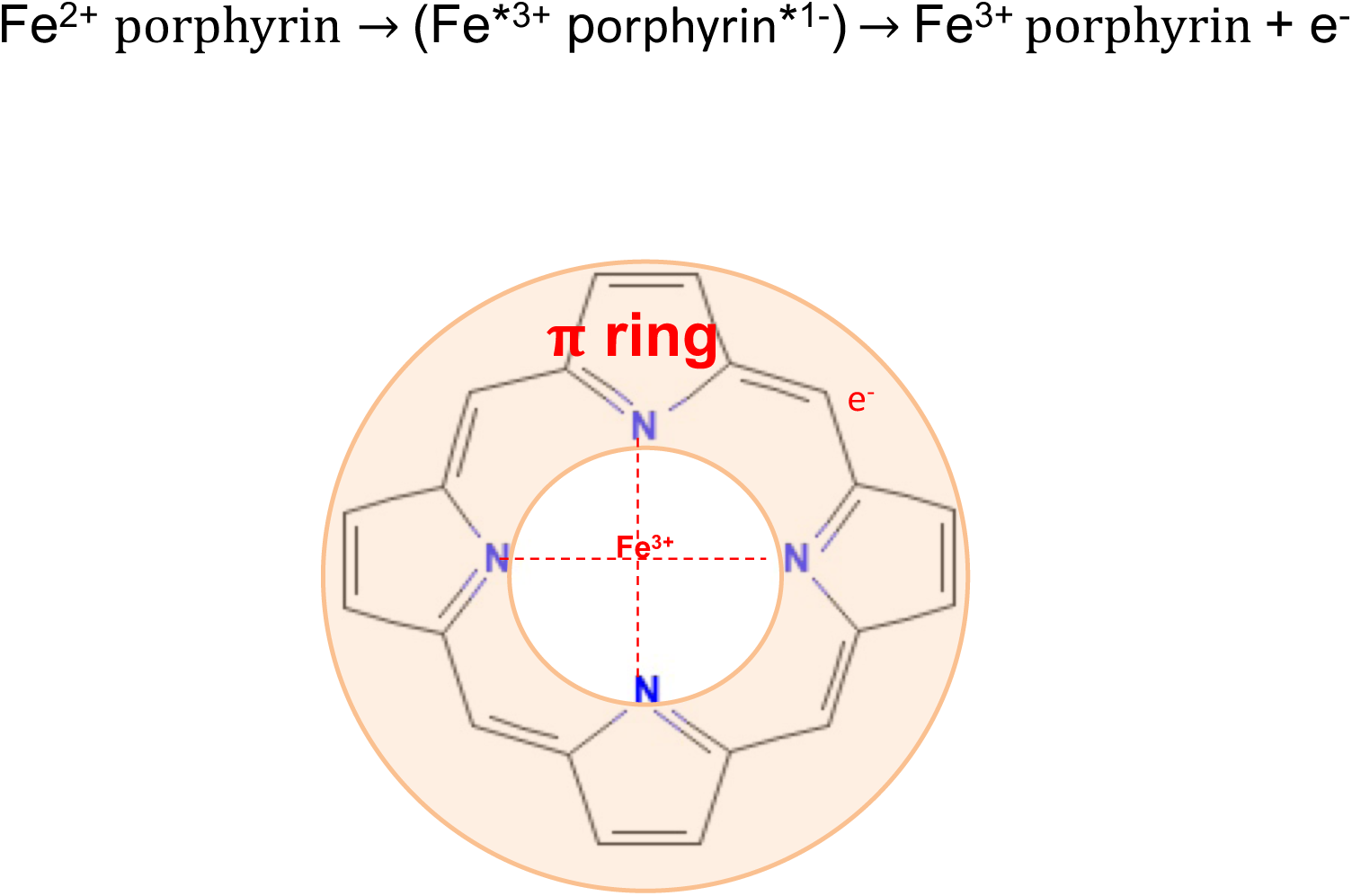
The potential mechanism for Complex II and III to form radical pair intermediate. The triplet state of Fe*^3+^-*π*^*1-^ can form a stabilized radical pair intermediate state in the Fe^2+^-porphyrin system.

As for Complex V, its link to catalytical reactions could form charge transfer radical pair intermediates during the catalytical process. The catalytical reaction involves electron transfer between two molecules, temporarily generating spatially separated radical pair intermediates. Complex V produces ATP in the F_1_ region of the ATP synthase via ADP + Pi + H^+^ <= ATP + H_2_O, where Pi is an inorganic phosphate (Buchachenko, Kuznetsov et al. 2004, Buchachenko, Kouznetsov et al. 2005, Buchachenko, Kuznetsov et al. 2006). Mg^2+^, a singlet ground state with two paired electrons, plays an essential catalytic role in driving this ADP reaction (Igamberdiev and Kleczkowski 2015) by coordinating itself with the nearby oxygen atom of ADP (Gout, Rebeille et al. 2014). Buchachenko et al. (Buchachenko, Kuznetsov et al. 2006) have proposed that a radical pair of Mg^•1+^-ADP^•2-^ intermediate is formed by detaching one electron from oxygen in ADP^3-^ to Mg^2+^, which can be affected by LMF. To support this idea, they used ^25^Mg^2+^, the isotope of ^24,26^Mg^2+^, to generate an internal MF, which interacts with the two-electron singlet state of Mg^•1+^-ADP^•2-^, much the same as the radical pair mechanism, to induce LFE that enhanced ATP synthesis rate (Buchachenko, Kuznetsov et al. 2004, Buchachenko, Kouznetsov et al. 2005, Kuznetsov 2014). The near 40% Vmax increase, after adding ADP, suggests that Mg^•1+^-ADP^•2-^ radical pair intermediate may account for the Complex V LFE.

As mentioned above, the Complex I activity did not show an LFE. The curve fitting shows a trend of monotonical decreases in activity but not statistically significant with the LMF. Therefore, the iron-sulfur clusters in Complex I likely only support electron transfer but have no radical pair intermediate formation. Uniquely, Complex I starts its activity differently from Complex II, with a proton-coupled electron transfer between NADH and FMN (NADH + H^+^ + FMN → NAD^+^ + FMNH_2_) before transferring two electrons through seven iron-sulfur clusters (Hannah R. Bridges 2012) except N1a 2Fe-2S cluster. The two electrons are transferred one at a time. The first electron is transferred via NADH + FMN → NAD^+^ + FMNH* + e^−^, as suggested by Saura and Kaila (Saura and Kaila 2019), followed by H^+^ + FMNH* → FMNH_2_, and then transferred the first electron to the next iron-sulfur cluster. The generated intermediate FMNH* is critical in modulating the second electron transfer, hence the total Complex I activity. This intermediate has a single electron at a doublet spin state (D). The applied MF can split the doublet spin state into a higher energy D_+1/2_ and a lower energy D_-1/2_ spin state. With an increasing MF, the intermediate can populate more into the lower energy spin state D_-1/2_. Since the electron transition from NADH to FMNH_2_ has a relatively small reduction potential (Zu, Shannon et al. 2003, Treberg and Brand 2011, Holt, Efremov et al. 2016), the intermediate population at the lower D_-1/2_ may stabilize to hinder the transfer of the first electron to the iron-sulfur cluster in Complex I. Such a stabilization of the intermediate state population could decrease the Complex I activity by hindering the subsequent NADH binding to transfer the second electron. This may explain why we have observed a “slight” decreased Complex I activity trend by LMF.

Regarding the effects of LMF on ROS, most ROS are generated in Complexes I and III through electron leakage/reverse electron transfer (Brand 2016, Hernansanz-Agustin and Enriquez 2021). In the presence of an LMF, the forward electron transfer is enhanced via the LFE, consequently decreasing the reverse electron transfer to generate ROS. This is reflected in Figure 7B, which shows an inverted bell-shaped LFE effect on H_2_O_2_. Interestingly, such an effect is absent when Complex III is blocked by antimycin, suggesting that the LMF-mediated changes in ROS generation require Complex III to be functional.

The activities of TCA cycle enzymes citrate synthase and succinate dehydrogenase (Complex II) also show a bell-shaped LFE but not malate dehydrogenase (Figure 7). Succinate dehydrogenase plays a dual role in OXPHOS by catalyzing the oxidation of succinate to fumarate in the TCA cycle and converting FAD to FADH_2_ to initiate the Complex II activity.

Therefore, the above-described explanation for Complex II LFE applies to the response of succinate dehydrogenase to the LFE. Only two reactions in TCA cycles involve aconitase enzyme in mitochondria as a catalyst; both are downstream reactions to the citric synthase. Citrate synthase generates citrate, which is converted to isocitrate via the catalytical activity of aconitase. Aconitases are metalloenzymes containing iron-sulfur clusters. The active form of the enzyme contains a [4Fe–4S] cluster. Water molecules coordinate the labile iron atom (Feα) and are essential for the enzyme’s catalytic function. The Feα atom within active aconitase coordinates with citrate and water oxygen atoms. It acts as a Lewis acid to activate the hydroxyl group of citrate and facilitate the isomerization reaction. The Fe^2+^-citrate intermediate could delocalize one Fe^2+^ electron to citrate in this step (Castro, Tortora et al. 2019). This delocalization could potentially create a charge-separated radical pair intermediate of Fe^3+^- citrate^−^ to generate LFE. As for malate dehydrogenase, we have no reasonable explanation for its single positive response to the LMF. However, the better straight-line fitting of the data suggests it may not be from the LFE.

The discussion above indicates that cytochromes, iron-sulfur clusters, and catalytical reactions within the mitochondrial ETC and TCA cycle are responsible for the LFE in the observed results. We propose that radical pair intermediates are formed in these three sites, which interact with the LMF to evoke spin state transition from S_0_ to T_0_, leading to enhanced activity of OXPHOS for increasing mitochondrial bioenergetics. Intriguingly, our observed LFE on several OXPHOS enzymes all occurred around the same range of LMF, as was detected in many other systems (Karimi, Ghadiri Moghaddam et al. 2020). As is discussed in Supplementary Materials, the mechanism behind the LFE is primarily attributed to the suppression of hyperfine interactions by an applied LMF. These enzymes in mitochondria have a similar hyperfine energy range and thus respond to a similar range of LMF for achieving LFE.

### LMF mitigation on I-R injury

Our data show a bell-shaped increase in Vmax in rat mitochondria after I-R treatment. Such an increase in the ATP generation can ease the I-R injury level or recovery from it. In addition, the decrease in ROS on control mitochondria also pointed out a potential mechanism for decreasing I-R injury by LMF. During the I-R process, a sudden surge of ROS occurs after reintroducing O_2_. Succinate accumulates during cardiac ischemia, which drives electrons through Complex II to reduce CoQ to CoQH_2_ (Chouchani, Pell et al. 2014). Since the absence of O_2_ stalls the ETC, the electrons are driven back through Complex I to reduce FMN. On reperfusion with O_2_, the excess Complex I electrons are transferred from FMN directly to O_2_ to generate O_2_^●-^.

Three approaches could be used to mitigate this destructive I-R injury. The first approach could be to minimize the level of succinate in the heart mitochondria by increasing the transfer of succinate to fumarate, malate, and citrate, with the citrate exported from the mitochondrion by the citrate carrier (SLC25A1). The second approach could impair Complex I to minimize reverse electron transfer, FMN reduction, and mROS production. The third could be to reduce the mitochondrial inner membrane potential and ramp up the forward electron flux through the ETC, thus oxidizing CoQH_2_ and minimizing reverse electron transfer.

These three strategies seem achievable when cardiac mitochondria are exposed to a static LMF of ∼3 to 7×10^−4^ T. The specific activities of succinate dehydrogenase and citrate synthase show LFE, which would deplete mitochondrial succinate. The minor effects of LMF on the activity of Complex I suggest that it would not promote reverse electron transfer. The forward electron flux is increased by the activation of the ETC in association with the activation of the ATP synthase to depolarize the mitochondrial membrane potential. Finally, mROS drops to a minimum at ∼4 to 6×10^−4^ T. It would seem an extraordinary coincidence that this coordination of alterations in many mitochondrial functions could all occur by chance due to an increase in the LMF surrounding the mitochondrial of average ∼3 to 7×10^−4^ T.

### Conclusion/Summary

In summary, we have evaluated the effect of LMF, between 0 to ∼20×10^−4^ T, on the activities of mitochondrial ETC and TCA cycle proteins. RCR, Complexes II, III, and V, and citric synthase and succinate dehydrogenase show increased activities with similar bell-shaped LFE responses. Instead, Complex I shows no or a slight negative response trend to LMF. Moreover, the bell-shaped increases in RCR are also preserved in mitochondria after I-R injury.

Interestingly, ROS levels show an inverted bell-shaped LFE. To explain these effects, we have proposed theoretical mechanisms via intermediate radical pair formation and electron spin transition in cytochromes, iron-sulfur clusters, and catalytical reactions of magnesium-ADP.

This study raises several questions for future exploration: (a) Can the LFE in mitochondria observed in this study also couple to a longer-term redox signaling to influence various cellular processes like cell proliferation, differentiation, migration, and gene expression, as shown preciously? (b) Are there additional LMF-mediated processes, such as alteration of cristae architecture and curvature and location selective ETC supercomplexes formation, which can also influence mitochondrial bioenergetics? As the electrons transverse Complexes I, III, and IV, protons are pumped from the mitochondrial matrix into the closed cristae lumens, where they are sequestered (Pham 2016, Ren, Nemati et al. 2020). In intact cells, it has been shown that the cristae of adjacent mitochondria become aligned. The authors suggested that the concentrated protons may generate an endogenous electromagnetic field to promote such an alignment (Picard 2015). (c) What are the biological impacts of mitochondria exposure to zero MF environments, like astronauts in a spaceship? (d) Does the LMF have an intrinsic hormesis characteristic in biological systems where low intensities are beneficial and high intensities are detrimental? (e) Can we apply LMF to improve human health and disease by boosting cellular energetics while lowering oxidative stresses? (f) Can exposure of mitochondria to specific wavelengths of light augment the effect of LMF? The data presented here may help stimulate future studies addressing these questions.

## Materials and Methods

Animals: Ethics Procedures were in strict accordance with the Division of Laboratory Animal Medicine, University of Rochester, in compliance with state law, federal statute, and NIH policy, and were approved by the Institutional Animal Care and Use Committee of the University of Rochester (University Committee on Animal Resources (UCAR) protocol 101172/2011-008).

Male Sprague Dawley rats were obtained from Charles River (Wilmington, MA). Rats were anesthetized with CO_2,_ which was confirmed by tail pinching. The hearts were removed quickly and transferred into an ice-cold buffer or attached to a Langendorff perfusion system

### Material

Unless stated otherwise, all chemicals were purchased from Sigma (St. Louis, MO). Sprague Dawley rats were obtained from Charles River (Wilmington, MA).

### Isolation of heart mitochondria

Mitochondria from rat hearts were isolated, according to Sokolova et al.(Spinazzi, Casarin et al. 2012). Briefly, minced heart tissue was subjected to treatment with 5 mg subtilisin, which was dissolved in 10 ml of isolation medium A (225 mM mannitol, 70 mM sucrose, 0.5 mM EGTA, 0.5 mM EDTA, and 20 mM HEPES pH 7.4), for 8 minutes at room temperature while gently stirring. The protease reaction was stopped by adding a 10-fold excess of fatty acid-free bovine serum albumin dissolved in isolation medium A. The tissue was then homogenized with an Elvejhem potter, and the mitochondria were separated from the homogenate by differential centrifugation at 500 g and 9,000 g. The final mitochondrial sediment was resuspended in 225 mM mannitol, 70 mM sucrose, and 10 mM HEPES pH 7.4.

The protein concentration of the isolated mitochondria was about 18 mg/ml, as determined by Lowry. Mitochondria were kept on ice and used for functional assays within 3 hours after finishing the isolation protocol.

### Determination of respiratory control index (RCI)

Directly after completing the isolation protocol to obtain isolated heart mitochondria, we determined the respiratory control index (RCI) as a measure of mitochondrial functionality with a Clark-type oxygen electrode from Hansatech (PP Systems, Boston, MA). The measurements were carried out at room temperature in a 2 ml respiration medium containing 70 mM mannitol, 25 mM sucrose, 20 mM HEPES, 120 mM KCl, 5 mM KH_2_PO_4_, 3 mM MgCl_2,_ pH 7.4. Substrate-mediated respiration (V0) was initiated by adding 3 mM malate and 5 mM glutamate. Then, 1 mM ADP was added to obtain maximal respiration (Vmax). The RCI was calculated by dividing the slope of Vmax by Vo (Villani and Attardi 2001, Beutner, Sharma et al. 2005). At the end of each experiment, 8 μM cytochrome c and 10 μM atractyloside was added to test the intactness of the outer and inner mitochondrial membrane, respectively.

### Ischemia-reperfusion injury

Hearts from 8-10 weeks old male Sprague Dawley rats were retrogradely perfused in Langendorff mode under constant flow (12 ml/min) with 100% oxygenated Krebs–Henseleit buffer containing 118 mM NaCl, 4.7 mM KCl, 1.2 mM MgSO_4_, 24 mM NaHCO_3_, 1.2 mM KH_2_PO_4_, 2.5 mM CaCl_2_, 11 mM D-glucose, as described in (Nadtochiy, Tompkins et al. 2006). After 20 minutes of equilibration, hearts were subjected to 20 minutes of global ischemia by stopping the flow of Krebs-Henseleit solution perfusion without oxygenation, followed by 20 minutes of reperfusion with 100% oxygenated buffer solution.

### Activity tests for Complex I, II, III, and V

The enzymatic activities of the ETC Complexes were measured using the protocols described in Spinnazzi et al.(Spinazzi, Casarin et al. 2012).

Complex IV electron transport involves copper centers and is not investigated in our studies. All tests were done at room temperature, and the activities were measured with a Genesys 5 UV/Vis spectrophotometer (Thermo Scientific, Pittsburgh, PA). Per test volume of 1 ml, 5-20 μg of mitochondrial protein was used. Mitochondria were frozen and thawed at least once to assess the activity of the complexes, but not more than 3 times to avoid a substantial decrease in activity due to protein degradation.

The NADH-ubiquinone oxidoreductase activity of Complex I was measured in an assay buffer (50 mM potassium phosphate pH 7.6, 1 mM EDTA, 2.5 mg/ml fatty acid-free bovine serum albumin (Roche Bioscience), 2 mM potassium cyanide, 1 μM antimycin A and 0.1 mM NADH) and started by adding 65 μM ubiquinone_1_ to the assay mix. The decrease of NADH was followed at 340 nm, and the activity was calculated using an extinction coefficient (ε) for NADH of 6.81 mM^−1^cm^−1^. Each test was repeated with rotenone (2μg/ml) to obtain the complex I-specific, rotenone-sensitive activity.

Complex II activity was measured as succinate: ubiquinone_1_ oxidoreductase, where the generation of FADH was utilized by the artificial electron acceptor 2,6-dichlorophenolindophenol (DCPIP, ε = 19.1 mM^−1^cm^−1^). The absorbance was followed at 600 nm for 3-5 minutes in an assay buffer (25 mM potassium phosphate pH 7.2, 5 mM MgCl_2_, 20 mM sodium succinate, 2 mM potassium cyanide, 50 μM DCPIP, 2 μg/ml antimycin A and 2 μl/ml rotenone). The reaction was started by adding 65 μM ubiquinone_1_.

The activity of Complex III was accessed as succinate-cytochrome c reductase in an assay buffer (25 mM potassium phosphate pH 7.2, 5 mM MgCl_2_, 2.5 mg/ml BSA, 2 mM potassium cyanide, 2μg/ml rotenone, 0.6 mM n-dodecyl-maltoside, 35 μM ubiquinol_2_ and 15 μM oxidized cytochrome c (to start the reaction)). Each test was repeated in the presence of 1 μM antimycin A to account for antimycin A-insensitive background activities. The reduction of cytochrome c was measured at 550 nm (ε = 18.7 mM^−1^ × cm^−1^).

The activity of Complex V was assayed as F_1_-ATPase. This assay links the ATPase activity to NADH oxidation via the conversion of phosphoenolpyruvate to pyruvate by the pyruvate kinase. Each test was repeated in the presence of 2 μM oligomycin to account for oligomycin-insensitive ATPase activity.

### Activity tests for selected TCA cycle enzymes

These tests were done according to protocols published by Reisch and Elpeleg (Reisch and Elpeleg 2007). All tests were done at room temperature.

The malate dehydrogenase catalyzes the reversible conversion of oxaloacetate into malate in the presence of NADH. The generation of NAD^+^ was measured at 340 nm in an assay buffer (50 mM potassium phosphate, pH 7.4, 2 mM NADH, and 0.01 % Triton X-100). The reaction was started by adding 0.5 mM oxaloacetate.

Citrate synthase activity was followed using a linked assay by condensing acetyl CoA and oxaloacetate to form citrate and CoA-SH. The latter reacts with 5,5-dithiobis-(2-nitrobenzoic acid) to form 5-thio-2-nitrobenzoate, which has a yellow color and is monitored by the increase of absorbance at 412 nm (Kirby, Thorburn et al. 2007).

Succinate dehydrogenase was measured using a variation of the above-mentioned Complex II activity assay. This test measures the conversion of succinate to fumarate, and phenazine methosulfate is used as an artificial electron acceptor to reduce cytochrome c. Potassium cyanide (2 mM) is included to prevent reoxidation of cytochrome c by Complex IV. The reaction was started by adding succinate and the change of absorbance was measured at 550 nm and ε is 19 mM^−1^ × min^−1^.

### Detection of ROS

ROS were detected using the Ampliflu Red^TM^ fluorescent dye (10-acetyl-3,7- dihydroxyphenoxazine from Sigma St. Louis IN), which reacts with H_2_O_2_ in the presence of horseradish peroxidase in a 1:1 stoichiometry, producing highly fluorescent resorufin.

Superoxide dismutase was added to the test to convert O^2*-^ into H_2_O_2_. For these experiments, 1 to 1.5 mg mitochondrial protein was suspended in 2 ml respiration medium (70 mM mannitol, 25 mM sucrose, 20 mM HEPES, 120 mM KCl, 5 mM KH_2_PO_4_, 3 mM MgCl_2,_ pH 7.4) and exposed to the indicated LMF for 2 minutes. ROS were measured immediately with a Cary Eclipse fluorescence spectrophotometer (Varian Inc. Walnut Creek CA) using an excitation wavelength of 563 nm, and the emitted fluorescence was detected at 587 nm. V_0_ was induced by adding 5 mM glutamate and 3 mM malate. Vmax was induced by adding 5 mM glutamate, 3 mM malate, and 1 mM ADP. Then, the Complex III inhibitor antimycin A (2 µg/ml) was added to maximize ROS generation. At the end of each experiment, defined concentrations of H_2_O_2_ were added to the cuvette to calibrate the signal.

### Statistical Analysis

All values are expressed as mean ± SE. Data were analyzed Graphpad Prizm v9.5.1, and p≤0.05 was determined to be statistically significant. Non-linear regression fittings of the data vs. the applied MF were performed with Graphpad Prizm v9.5.1. A second-order polynomial correlation was tested against a first or third-order alternative hypothesis. R^2^ and p values were used to determine the agreement and statistical significance of the curve fitting.

## Acknowledgement

This research was triggered by a hallway conversation between Arthur J. Moss (AJM) and Shey-Sing Sheu (SSS) in 2010 at URMC. AJM was a cardiologist caring for many patients with implanted pacemakers. He asked SSS why the heart tissue adjacent to pacemakers appears to become healthier after implantation. SSS replied, hypothetically, that it could be due to the beneficial effects on mitochondrial bioenergetics and redox signaling by the small electromagnetic field originating from pacemakers. They both thought this was an interesting and testable hypothesis and thus launched the investigation subsequently. The setup of the Helmholtz coil in the measurement chamber of oxygraph was in consultation with Professor Mark Bocko (Department of Electrical and Computer Engineering, UR). Most experiments were carried out by Gisela Beutner (GB) during 2011-2012 at URMC. The results were used to file a US patent granted in 2018 (Low Intensity Magnetic Field Devices for Treatment of Cardiac and Neurological Disorders, Patent No: US 9,873,000 B2). The main reasons for the lengthy delay in publishing these results were 1) we were not able to interpret the surprising bell-shaped LFE of our results until the participation of Huoy-Jen Yuh (HJU) in 2021, who proposed the theoretical radical-pair mechanisms, and 2) relocation of SSS to TJU in 2011. We want to thank all those who have contributed to this work through critical discussion and constructive ideas.

Specifically, we want to thank the volunteers of Goldenberg and Emily Rowland, who participated in some of the described experiments. Finally, AJM passed away on February 14, 2018. We would like to dedicate this manuscript to memorizing AJM’s legacy as a physician, scientist, teacher, and mentor.

## Supplementary figure legends

**Supplementary Figure 1:**
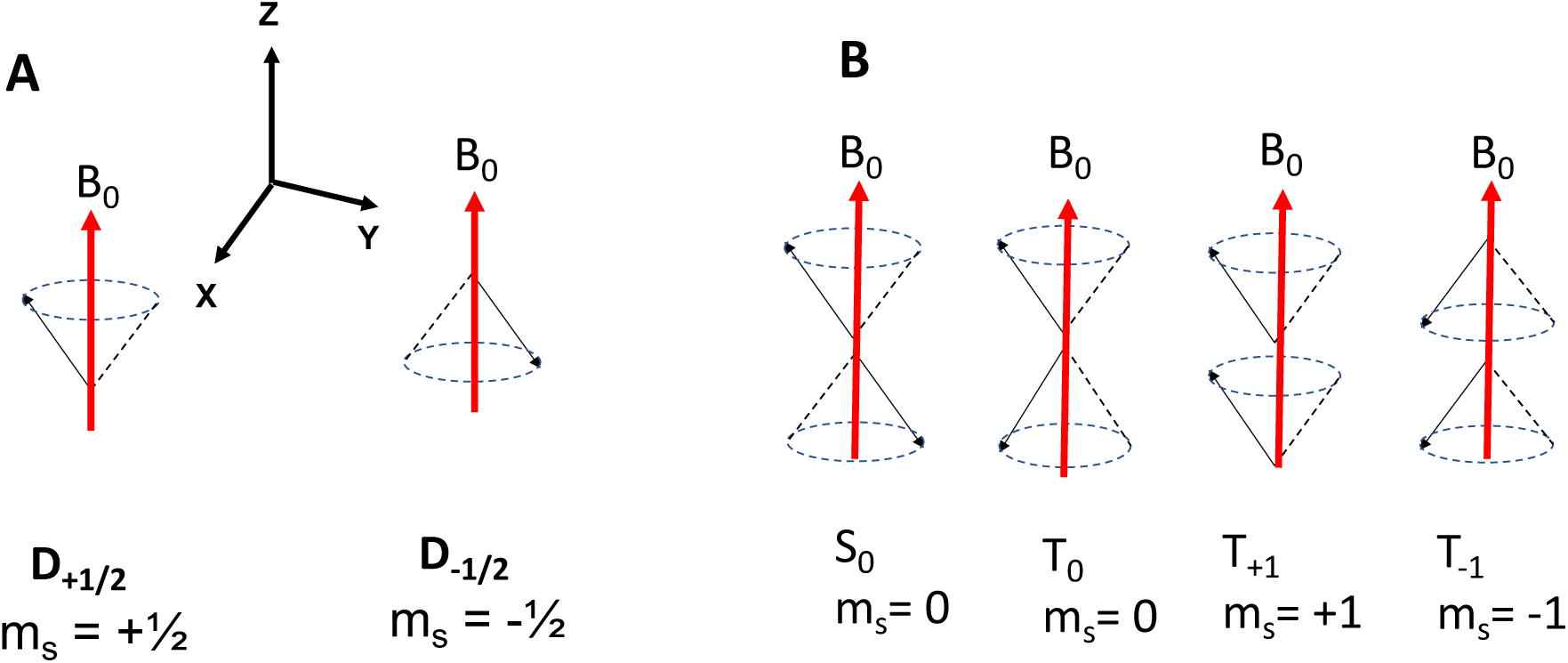
Classic vector representation of electron spin moment (m_s_) precession along the MF (B_0_), defined as the z-axis. **A:** Spin vector precession of one electron spin, D_1/2_, along the MF (red arrow) with upward, m_s_ = +½, and downward, m_s_ = -½, direction**. B:** Spin vector precession along the MF with paired two electron spins, S_0_, and unpaired two electron spins, T_0_, T_+1_, and T_-1_.

**Supplementary Figure 2:**
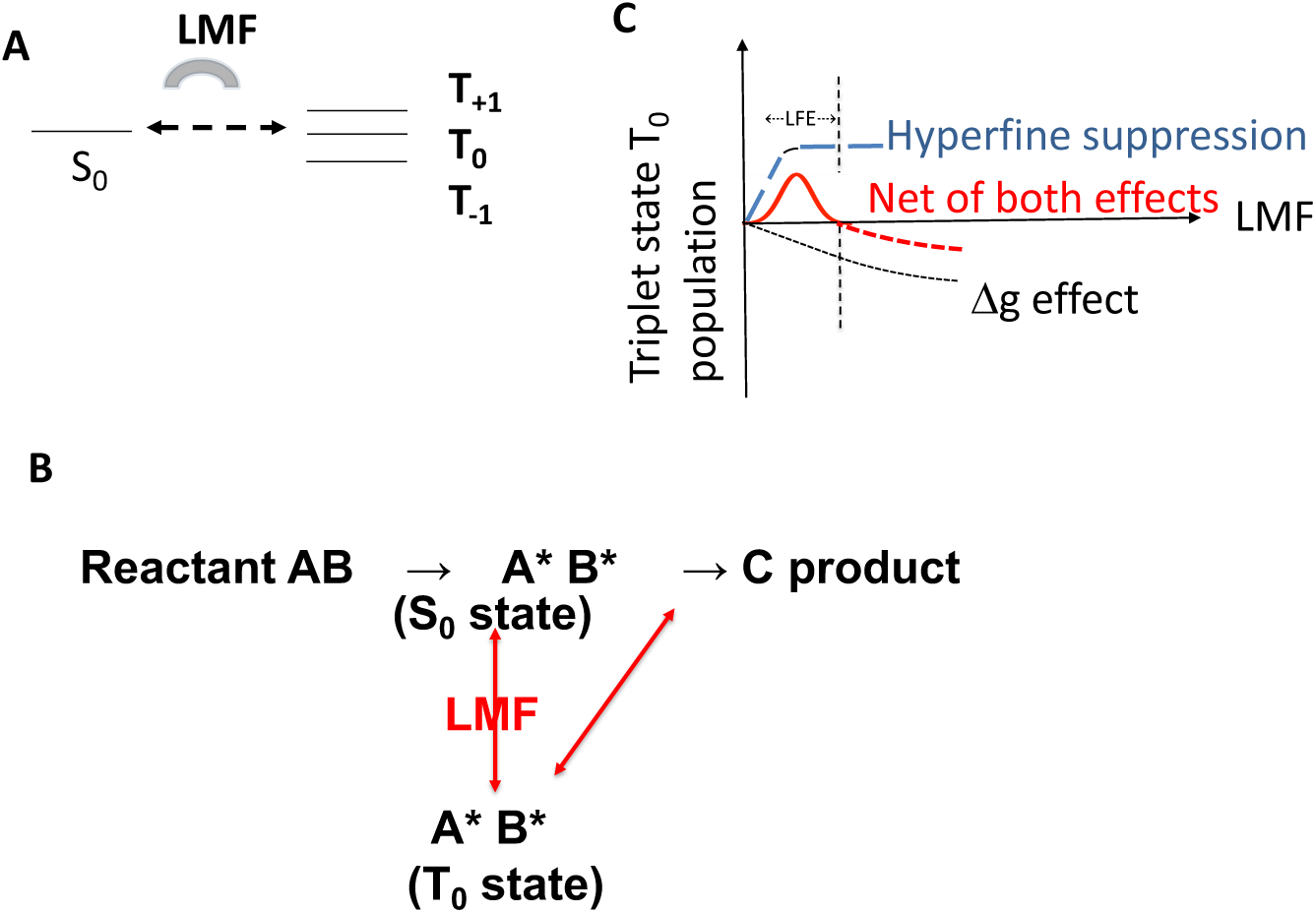
A radical pair mechanism of an LMF produces a low field effect (LFE) through the spin transition. A: Schematic chemical reaction diagram showing alternative reaction pathways among reactant AB, radical pair A*B* at S_0_ state, radical pair A*B* at T_0_ state, and product C. The LMF creates the spin transition that favors the reaction from A*B* at the T_0_ state to the product C. B: LMF affects the energy levels of the triplet states via the Zeeman effect that promotes the spin transition from S_0_ to the T_0_ state. C: Very LMF affects the singlet to triplet transition by increasing the S_0_ to T_0_ transition due to hyperfine degeneracy suppression (blue line, hyperfine suppression). As the MF increases, Δg effects become pronounced, which decreases the transition from S_0_ to T_0_ (black line). Combining both effects creates a bell-shaped population of T_0_ as a function of an LMF (red line).

## Additional information

### Author Contributions

G.B., A.J.M., and S-S.S. designed research; G.B. and H-J.Y. performed experiments; G.B. and H-J.Y. analyzed data; H-J.Y. and D.C.W. theoretical interpretation of data; I. G., A.J.M., D.C.W., and G.A.P. participated in research discussion; G.B., H-J.Y., D.C.W., G.A.P., and S-S.S. wrote the paper.

