## Supplementary Materials for "Low Magnetic Fields Stimulate Cardiac Mitochondrial Bioenergetics with a Bell-Shaped Response: Possibly Via a Radical Pair Mechanism"

To understand how a low magnetic field (LMF) can interact with the oxidative phosphorylation process, which involves the transport of electrons in mitochondria, it is worthwhile to briefly describe the quantum mechanics of LMF influences on electron magnetic spin moments and radical pair reaction.

**Spin Moment Interacts with a Magnetic Field**

According to quantum physics, only spins of particles, molecules, or atoms can directly interact with a magnetic field (MF) (Hayashi 2004, Eason 2019) by aligning themselves with the field. The observed magnetic effects result from the alterations of their spin states and energies by their interaction with an MF. In quantum mechanics, electron spin has its intrinsic spin angular momentum, represented by the spin quantum number "S." An electron has a total 2S+1 number of spin states, with each state corresponding to a spin moment m_s_, also known as the spin magnetic quantum number. m_s_ defines its spin angular momentum and direction relative to an applied MF. A single electron has a spin quantum number S of ½ and has two spin states with their associated m_s_, +½ for aligning with the field (“spin up”) and -½ for aligning against the field (“spin down”). In a classic model, the spin moment of one electron processes in a cone shape along the MF with either up or down directions to the field, as shown in Supplementary Figure 1A. The electronic spin states of an atom or a molecule depend on the net spin of their total electrons. For those with two paired electrons that have opposite spin directions to each other, their spins cancel each other out, resulting in a net spin S being 0 (S_0_), a singlet state, with only one spin state of m_s_ = 0 (Supplementary Figure 1B). For those with two unpaired electrons, the net spin S is 1, with three spin states of m_s_ being 0, +1, and -1 (Supplementary Figure 1B). In other words, an atom or molecule’s "net spin" refers to the total number of unpaired electrons (n) divided by 2, n/2. The atoms or molecules with n number of unpaired electrons have spin S= n/2 and 2S+1 = n+1 possible spin states. For example, a molecule with one single electron, where the net spin S is ½, has two spin states (a doublet state), D_+1/2_ and D_-1/2_, with m_s_= +½ and -½ (Supplementary Figure 1A). For two unpaired electrons, the net spin S is 1 with three spin states, triplet states of T_+1_, T_0_, and T_-1,_ where the net spins are parallel (m_s_= +1), perpendicular (m_s_= 0), or antiparallel (m_s_= -1) to the quantum axis (Supplementary Figure 1B). The two electron spins in S_0_ and T_0_ are in opposite directions but in phase in S_0_ and out of phase in T_0_, as shown in Supplementary Figure 1B. All triplet states have spin angular moments on the xy plane, but not those in singlet states. In the presence of an MF, non-zero electron spins can interact with MF, known as the Zeeman effect, to alter its energy, ΔE=*g*β_e_B_0_m_s_/*h*, where *g* is the g-factor, β_e_ is the Bohr magneton, B_0_ is the MF, and *h* is Planck constant. Such an interaction splits spin energy among different m_s_, with spin energy increasing with positive m_s_, no change with zero m_s_, and decreasing with negative m_s_. The magnitude of energy splitting increases with increasing MF. The MF also alters their Larmor precession speed. In the classic vector precession model, the interaction of electron spin with an MF will cause the electron's magnetic moment vector to precession around the direction of the applied MF with a Larmor frequency =*g*β_e_B_0_/*h*. This precession frequency, changed by the MF, can affect the interconversion between different spin states.

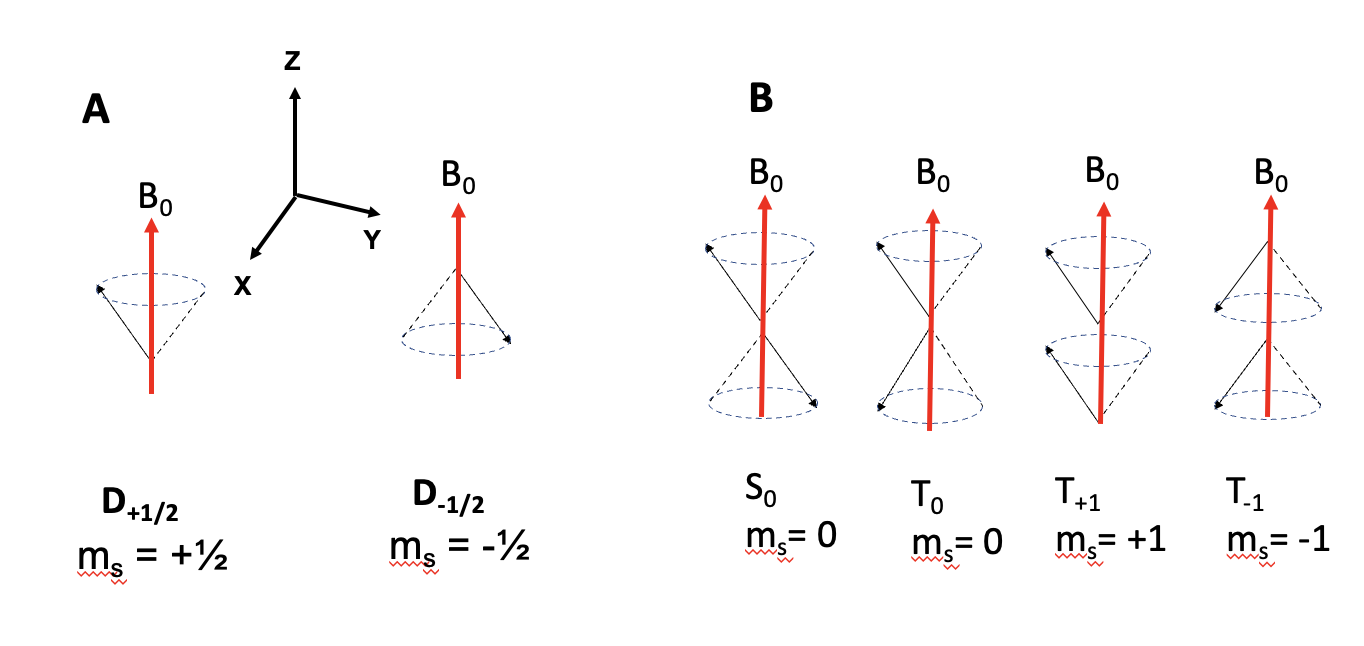

**Supplementary Figure 1:** Classic vector representation of electron spin moment (m_s_) precession along the MF (B_0_), defined as the z-axis. **A:** Spin vector precession of one electron spin, D_1/2_, along the MF (red arrow) with upward, m_s_ = +½, and downward, m_s_ = -½, direction**. B:** Spin vector precession along the MF with paired two electron spins, S_0_, and unpaired two electron spins, T_0_, T_+1_, and T_-1_**.**

**Radical Pair Mechanism in Spin Chemistry**

Interestingly, the energy splitting from the Zeeman effect, at an MF of less than 10^-3^ Tesla where Low Field Effect (LFE) is detected, is several orders of magnitude less than the thermal energy of kT, ~2-3x10^-2^ ev (electron volt) at room temperature. Such an energy shift from Zeeman interaction is too tiny thermodynamically to effectively account for the population change of the spin states in most chemical reactions. Therefore, other mechanisms involving kinetic effect must be considered for generating such a significant LFE. A “Radical Pair Mechanism” has been proposed to explain the LFE with a bell-shaped response to a weak MF of <10^-3^ Tesla (Y. Sakaguchi 1980, U Steiner 1989, Hore and Mouritsen 2016). Specifically, a chemical reaction responding to LFE involves an intermediate state formation of a “weakly” coupled “radical pair,” A*B*, from reactants A and B or a single reactant AB before proceeding to the final product C, as shown in Supplementary Figure 2A.

The radical pair mechanism proposes that the chemical reaction starts with reactants A and B or a molecule AB with two net electrons, either singlet or triplet states, proceeding to the intermediate “spin coupled” and “spatially separated” radical pairs of A*B*, each with a single electron. The short-lived radical pair is usually produced through decomposing reactants, electron transfer, or hydrogen transfer between reactants A and B or within a reactant AB. The spin-coupled radical pair forms 4 spin states, one paired singlet state, S_0,_ and 3 unpaired triplet states, T_+1_, T_0_, and T_-1_ (Supplementary Figure 2B). This differs from the usual two electron spins in an atom or a molecule. In a radical pair, each electron resides within a molecule in a different radical or spatial location, namely, the charge separation. Therefore, each electron spin is surrounded by a different chemical environment. Typically, the energy levels of the singlet and triplet states are different, and for a radical pair, it is a function of the radical pair distance. When the separation distance increases to around ≥ 1nm, the energy levels become comparable between the singlet and triplet states (Rodgers 2009, Hore and Mouritsen 2016). Under this condition, the singlet and triplet states can interconvert with each other via their “hyperfine interaction,” resulting from the spin moment of the electrons interacting with the magnetic moment of the nucleus (generated by spins of protons and neutrons) in the molecule. The magnetic interaction between an unpaired electron and the nearby nuclei within the radicals, a Zeeman interaction, slightly splits the energy levels of T_+1_ and T_-1_. Such an interaction, either isotropic or anisotropic, generates local spin moments on the xy plane and allows for the interconversion among the singlet and triplet spin states of the radical pair to equal population among all 4 spin states, a hyperfine interaction as shown in Supplementary Figure 2B. This is an essential requirement for the radical pair to generate LFE, that the energy levels of their singlet and triplet states are comparable to enable interconversion of the 4 spin states. It makes the spin-coupled radical pair sensitive to external MF to alter their spin state populations. For a reactant AB starting with two “paired” electrons, an S_0_ state, the intermediate radical pair has 4 spin states, a singlet state and 3 triplet states. All 4 spin states of the radical pair can proceed to the final product C, as shown in Supplement Figure 2A. However, only the singlet radical pair can recombine with the original reactant, not the triplet radical pair. Hence, the triplet radical pair has a longer lifetime and a higher probability of yielding the product C. The radical pair mechanism enables an additional pathway for product generation. The applied external MF can alter the spin state population of singlet and triplet states at weak MF to alter the reaction product yield.

The equilibrium of the hyperfine 4 spin states population is disturbed by an external MF, B_0_, via Zeeman interaction, ΔE=*g*β_e_B_0_m_s_/*h* and Larmor precession frequency, *g*β_e_B_0_/*h*, as such, the populations of the singlet and triplet states are altered. This disturbance induces a bell-shaped change in S_0_ and T_0_ populations: it increases the transition between S_0_ to T_0_ at a weak MF and reverses the trend with further increasing MF strength. This bell-shaped effect has been explained by dividing it into two parts, as described by Sakaguchi et al. (Nagakura 1978, Y. Sakaguchi 1980) and Yiteng Zhang et al (Y. Zhang 2015). The first part is the hyperfine degeneracy disturbance (hyperfine suppression), and the second part results from the Δ*g*, *g* difference of the two radicals. When an MF is applied, starting at a very low value, the electron spins experience both the internal MF generated by nearby nuclei and the external MF. When the external MF is within the range of the hyperfine interaction energy, typical ~10^-6^ ev for molecules, around 10^-3^ T (Hayashi 2004), the radical pair precession along the combined field of both MFs. Such interaction alters the spin directions and their Larmor precession frequencies of the two radical spins. It increases the transition from singlet to triplet states. This increase in spin state transition mainly is from S_0_ to T_0_, as no flipping of spin direction is required (Hore and Mouritsen 2016). As the external MF suppresses the hyperfine interaction, the T_0_ population increases with increasing MF until saturation, as shown in Figure 2C (U Steiner 1989).

The external MF B_0_ split the triplet states' energy levels by increasing the T_+1_ energy level and decreasing that of T_-1_, but there were no changes in the S_0_ and T_0_. Such a splitting reduces the transition from the S_0_ and T_0_ states to T_+1_ and T_-1_ states while increasing the transition between the S_0_ and T_0_ states. As the MF increases, the split energy difference becomes more significant, causing the transitions of the S_0_ and T_0_ states to T_+1_ and T_-1_ to become less and less possible, resulting in only S_0_ and T_0_ interconversion remaining. As the MF increases, the internal MFs generated by the nuclei become insignificant, and the electron spins align mainly with the external MF direction. At this point, the second part, the *g*-factor differences, Δ*g*, between the electrons in two radical pairs, increasingly become more significant in altering their spin state population. The *g-*factor values of the two intermediate radicals are slightly different as they are a function of material-dependent interactions, characterizing their spin and orbital angular momenta. This g value difference, though small, can cause noticeable differences in Larmor precession frequencies of the two electron spins as the MF intensity gets higher. The increasing difference in their Larmor frequencies reduces the transition from S_0_ to T_0_, causing a decrease in the T_0_ population*.* It monotonically reduces the T_0_ state population until it reaches a saturation point at a much higher MF than that, causing hyperfine suppression (described in the first part), as shown in Supplementary Figure 2C. Combining both effects creates a bell-shaped population of T_0_ as a function of a weak MF, peaks less than 10^-3^ T. Multiple theoretical results have shown that the T_0_ state has a lifetime ~10^4^-fold longer than the S_0_ state (Lewis, Fay et al. 2018, Kerpal, Richert et al. 2019). This much longer lifetime of the triplet T_0_ state is the crucial reason why A*B* in the T_0_ state has a much higher probability of being the one to proceed to the final product C and, as such, induces LFE.

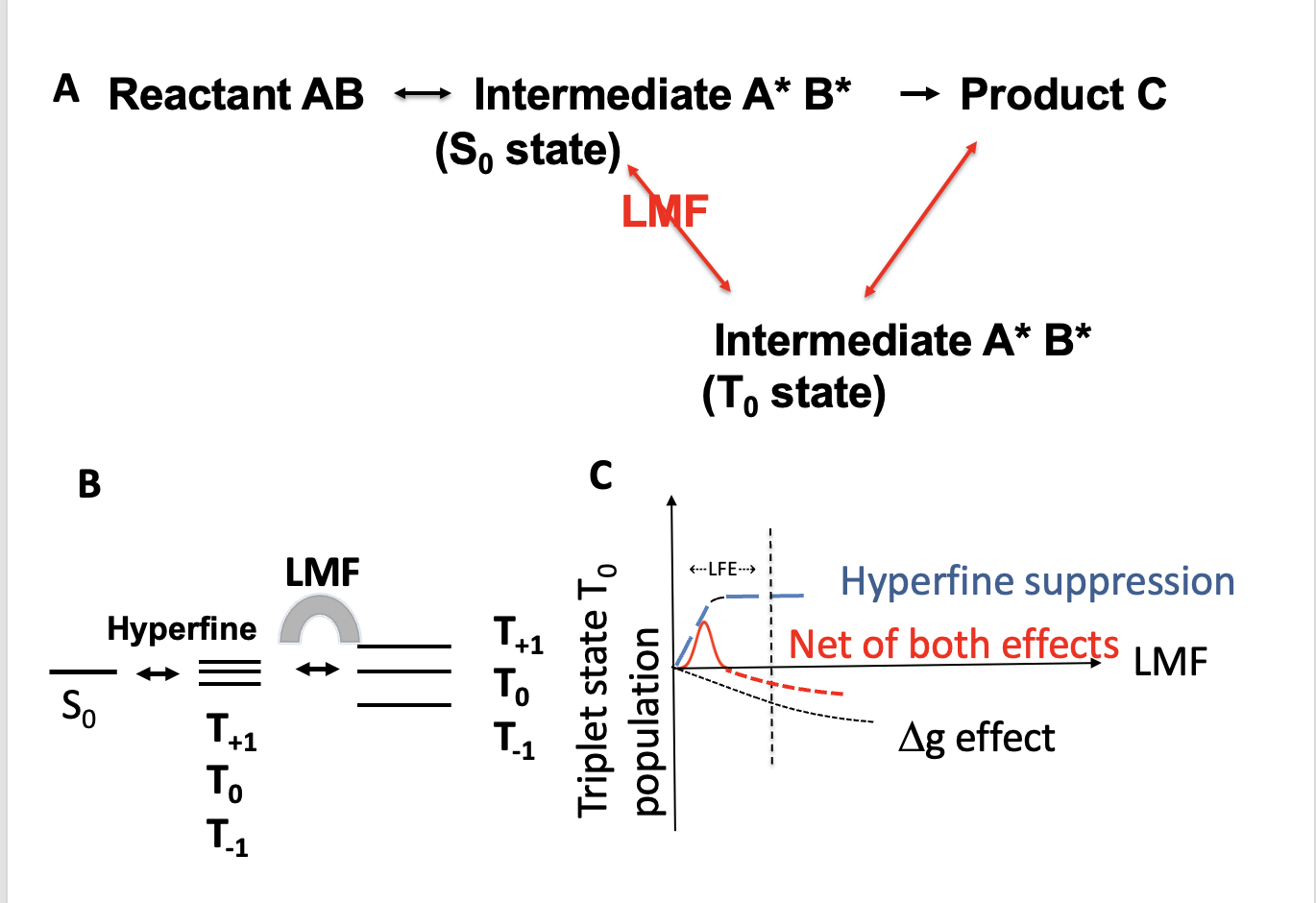

**Supplementary Figure 2:** A radical pair mechanism of an LMF produces a low field effect (LFE) through the spin transition. **A:** Schematic chemical reaction diagram showing alternative reaction pathways among reactant AB, radical pair A*B* at S_0_ state, radical pair A*B* at T_0_ state, and product C. The LMF creates the spin transition that favors the reaction from A*B* at the T_0_ state to the product C. **B:** LMF affects the energy levels of the triplet states via the Zeeman effect that promotes the spin transition from S_0_ to the T_0_ state. **C:** Very LMF affects the singlet to triplet transition by increasing the S_0_ to T_0_ transition due to hyperfine degeneracy suppression (blue line, hyperfine suppression). As the MF increases, Δg effects become pronounced, which decreases the transition from S_0_ to T_0_ (black line). Combining both effects creates a bell-shaped population of T_0_ as a function of an LMF (red line).

**Applying an external MF in the presence of an earth MF**

Though small, earth MF of ~5x10^-5^ T can still alter the radical pair hyperfine interaction to generate LFE. Recent studies on avian migration have generated interesting insights into the roles of the radical pair mechanism in birds’ compass magnetoreception (Hore and Mouritsen 2016, Xu, Jarocha et al. 2021). At this MF level, the Larmor precession frequency from the earth MF is 1.4 MHz, about 0.7 µs (microsecond) per period. The maximum possible MF effect can be predicted if the applied external MF generates a triplet state lifetime that exceeds the Larmor period (0.7 µs) (Hore and Mouritsen 2016). Such a singlet and triplet interconversion in a charge-separated radical pair, termed quantum beats, has been detected recently via the pump-push spectroscopy method (Mims, Herpich et al. 2021). The MF from 2.7 x 10^-4^ to ~1.5 x 10^-3^ T was applied in our experiments, which is 5-30 times higher than the earth MF, to generate LFE in OXPHOS enzymes in cardiac mitochondria. Future experiments to apply the spectroscopy method to detect the spin transition in mitochondrial proteins will validate our theoretical expalnations.

**115**(19): 1327-1341.
